# Evolution of PqsE as a *Pseudomonas aeruginosa*-specific regulator of LuxR-type receptors: insights from *Pseudomonas* and *Burkholderia*

**DOI:** 10.1101/2024.12.09.627592

**Authors:** Caleb P. Mallery, Kayla A. Simanek, Autumn N. Pope, Jon E. Paczkowski

**Affiliations:** Department of Biomedical Sciences, University at Albany, College of Integrated Health Sciences, Albany, New York, USA; Division of Genetics, Wadsworth Center, New York State Department of Health, Albany, New York, USA

**Author notes:** Address correspondence to Jon E. Paczkowski.

## Abstract

*Pseudomonas aeruginosa* is a Gram-negative opportunistic pathogen that poses a significant public health threat, particularly in healthcare settings. A key determinant of *P. aeruginosa* virulence is the regulated synthesis and release of extracellular products, which is controlled by a cell density-dependent signaling system known as quorum sensing (QS). *P. aeruginosa* uses a complex QS network, including two systems that rely on diffusible N-acylhomoserine lactone (AHL) signal molecules. The LuxR-type receptor RhlR is unique in that it requires not only its cognate AHL but also the accessory protein PqsE to maximally bind to promoter DNA and to initiate transcription. Our group demonstrated that PqsE physically interacts with RhlR, enhancing its affinity for target promoters across the *P. aeruginosa* genome. Although LuxR-type receptors are widespread in Gram-negative bacteria and important for pathogenesis, PqsE orthologs are restricted to *Pseudomonas* and *Burkholderia* species. This study explored the conservation of PqsE and examined PqsE ortholog structure-function across different species. Our results show that PqsE in *Pseudomonas* retain their functional interactions with RhlR homologs, unlike PqsE orthologs in *Burkholderia* spp., which do not interact with their respective LuxR-type receptors. Additionally, we assessed the AHL preferences of different receptors and hypothesized that the PqsE-RhlR interaction evolved to stabilize the inherently unstable RhlR, preventing its degradation. Indeed, we observe higher levels of RhlR protein turnover in a strain lacking *pqsE* compared to WT, which can be rescued in a strain lacking the Lon protease.

**IMPORTANCE:** *Pseudomonas aeruginosa*, a major pathogen for patients with cystic fibrosis and a primary constituent of healthcare-associated infections, relies on a complex quorum-sensing (QS) network to coordinate virulence factor production. Central to this system is the interaction between two proteins, PqsE and RhlR, which drive gene expression essential for pathogenesis. Our study investigates the conservation of the PqsE-RhlR interaction across related bacterial species, revealing that PqsE in *Pseudomonas* can enhance RhlR activity, while orthologs in *Burkholderia* lack this capacity. These findings offer new insights into the specificity and evolution of QS mechanisms, highlighting the PqsE-RhlR interaction as a potentially selective target for treating *P. aeruginosa* infections.

## INTRODUCTION

*Pseudomonas aeruginosa* is a Gram-negative pathogen found in a wide range of environmental niches, such as soil and water, that poses a significant public health threat, particularly in healthcare settings where it can colonize abiotic surfaces and in-dwelling medical devices^1^. Owing to its multidrug resistance and the production of extracellular virulence factors, secondary metabolites, toxins and biofilms, *P. aeruginosa* infections are difficult to treat and life-threatening^2^. *P. aeruginosa* is known to cause acute and chronic infections, which are prevalent in the lungs of patients with cystic fibrosis (CF)^3–5^. Individuals with CF, characterized by an inherited mutation in the CF transmembrane conductance (CFTR) gene, experience inadequate mucosal clearance due to the loss or partial loss of function of the chloride channel^6^. This deficiency results in the buildup of a nutrient-rich environment that fosters the growth of various bacteria. Historically, pulmonary infections during the early years of a CF patient’s life are caused by *Staphylococcus aureus* and *Haemophilus influenzae*^7^. As the disease progresses, *P. aeruginosa* emerges as the predominant pathogen, often succeeded by co-colonization with *Burkholderia cepacia* resulting in increased disease severity^8,9^.

One of the key determinants of *P. aeruginosa* virulence is the tightly regulated synthesis and release of extracellular products orchestrated by a cell density-dependent signaling system called quorum sensing (QS)^10–11^. QS is a cell-cell communication mechanism that relies on the production, detection, and community-wide response to autoinducer (AI) signaling molecules. *P. aeruginosa* utilizes three QS systems, two of which rely on diffusible *N*-acylhomoserine lactone (AHL) signal molecules^12^. AHL production and signaling are regulated by two conserved protein families, LuxI synthases and LuxR-type receptors: LuxI synthases catalyze the transfer of an acyl chain from an acyl-carrier protein (ACP) to *S*-adenosyl-L-methionine (SAM), producing an intermediate that undergoes lactonization to produce the final AHL product^12,13^. LuxI-produced AHL binds to LuxR-type transcription factor receptors possessing a variable N-terminal ligand-binding domain (LBD) and a conserved C-terminal helix-turn-helix (HTH) DNA-binding domain (DBD). AHL binding often stabilizes the receptor, allowing it to bind to DNA and regulate gene expression^14,15^. AHL production and detection are linked in a positive feedback manner, as LuxR-type receptors binding to their cognate AHL upregulate the expression of *luxR* and *luxI* genes, enhancing QS signaling pathways. In *P. aeruginosa*, two AHL systems (*las* and *rhl*) LasR and RhlR detect *N*-(3-oxododecanoyl)-L-homoserine lactone (3OC_12_HSL) and *N*-butyryl-L-homoserine lactone (C_4_HSL), respectively, which are produced by the LuxI synthases LasI and RhlI^16–18^. The third system, the Pseudomonas quinolone system (PQS), utilizes alkyl-4-quinolones (AQs), one of more than 50 produced by *P. aeruginosa*^19–20^. The *pqsABCDE* and *pqsH* genes encode the biosynthetic enzymes responsible for the synthesis of 2-heptyl-3-hydroxy-4-quinolone (PQS) and 2-heptyl-1H-4-quinolone (HHQ), which are chemically distinct from AHL signal molecules^21–23^. Both AQs can be detected by the LysR-family receptor PqsR (MvfR) to initiate the third wave of QS signaling in a mechanistically similar manner to AHL-dependent signaling. PqsR controls the expression of *pqsABCDE*, resulting in a positive feedback loop^24,25^. The final gene in the *pqsABCDE* operon, *pqsE*, encodes a thioesterase that hydrolyzes 2-aminobenzoylacetyl-coenzyme A (2-ABA-CoA) to 2-ABA in the PQS pathway but is dispensable for PQS production *in vivo*^26^. Our group and others discovered that the main function of PqsE is to regulate RhlR-dependent transcription through a physical interaction with RhlR, which enhances the affinity of RhlR for promoter DNA^27–30^. Indeed, RhlR, C_4_HSL, and PqsE coordinate the production of pyocyanin, a virulence factor that is often detected in the lungs of patients with CF^31^. Our group recently showed that PqsE physically interacts with RhlR, enhancing its affinity for target promoters, which is a previously unrecognized mechanism of QS regulation^27^. We also found that maximal RhlR-dependent transcriptional activation depends on C_4_HSL, the cognate AI of RhlR, across all binding sites^29^. Notably, PqsE and C_4_HSL exert different levels of influence on RhlR promoter binding and gene regulation. Therefore, there are two classes of RhlR promoters: C_4_HSL-dependent and PqsE-dependent^29^. This work established a hierarchical mode of RhlR-dependent gene expression and challenges the prevailing notion that RhlR and other regulators simultaneously activate their target genes^32^. Thus, RhlR and other LuxR-type receptors can have three distinct modes of activation: 1) ligand-dependent binding and gene expression, 2) ligand- and accessory protein-dependent binding and gene expression, and 3) ligand-independent and accessory protein-dependent binding and gene expression.

In this study, we investigated the evolutionary and functional conservation of the quorum-sensing accessory protein PqsE in *P. aeruginosa* compared to its orthologs, HhqE, in *Burkholderia* spp. Our findings reveal that PqsE, uniquely conserved in *P. aeruginosa*, forms dimers that enable its interaction with the LuxR-type receptor RhlR, stabilizing the receptor and enhancing its transcriptional activity. This interaction is absent in *Burkholderia* spp. HhqE orthologs, which lack key residues for dimerization and fail to complement PqsE function in the production of the RhlR-dependent virulence factors. We hypothesize that this necessary function evolved for PqsE and RhlR because RhlR is relatively insoluble in the presence of AI. Consistent with this, we show that RhlR is degraded by the Lon protease in the absence of *pqsE*, establishing for the first time the protective function of the evolved PqsE-RhlR interaction and providing new insights into the role of protein turnover in regulating QS signal progression.

## RESULTS

### Distribution of PqsE-RhlR proteins in Gram-negative bacteria

To determine whether the PqsE-RhlR interaction is a conserved mechanism of gene regulation, we first performed a targeted search to identify PqsE orthologs among Gram-negative bacteria. Our search of complete *Pseudomonas* genomes (https://www.pseudomonas.com)^33^ revealed that PqsE is restricted to *P. aeruginosa*, comprising 727 of the 728 *Pseudomonas* species (Table S1). We note that one strain, *P*. *fluorescens* NCTC 10783 was initially annotated as containing a PqsE homolog but has since been reclassified as *P. aeruginosa*^34^. This respiratory tract isolate carries a single amino acid substitution (G284R) distal to the PqsE-RhlR interaction interface on PqsE and has a fully conserved RhlR protein sequence. We include this strain as a naturally occurring PqsE variant (PqsE*^Pf^*) in our assays. Remarkably, 650 out of 728 *P. aeruginosa* strains maintain 100% amino acid identity for PqsE, indicating high conservation. Even in the least conserved sequences, PqsE maintains 91% sequence identity, and mutations are both rare and infrequent in clinical isolates. This level of conservation underscores the functional importance of PqsE in *P. aeruginosa*. The *pqs* biosynthetic operon of *P. fluorescens* NCTC 10783 displays a high degree of sequence identity in the *pqsABCDE* operon in *P. aeruginosa* (Fig. 1a). Consistent with previous reports, a broader search of the complete and incomplete bacterial genomes across the NCBI protein database showed that PqsE orthologs are restricted to the genus *Burkholderia*. While broader searches outside of *Pseudomonas* and *Burkholderia* yielded additional hits, these are unlikely to be true PqsE orthologs. PqsE orthologs founds outside of these genera are not located within discrete biosynthetic gene operons. Furthermore, the metallo-β-lactamase fold is highly prevalent among unrelated enzymes^35^. Thus, these potential orthologs likely represent functionally distinct proteins.

**FIG 1.**
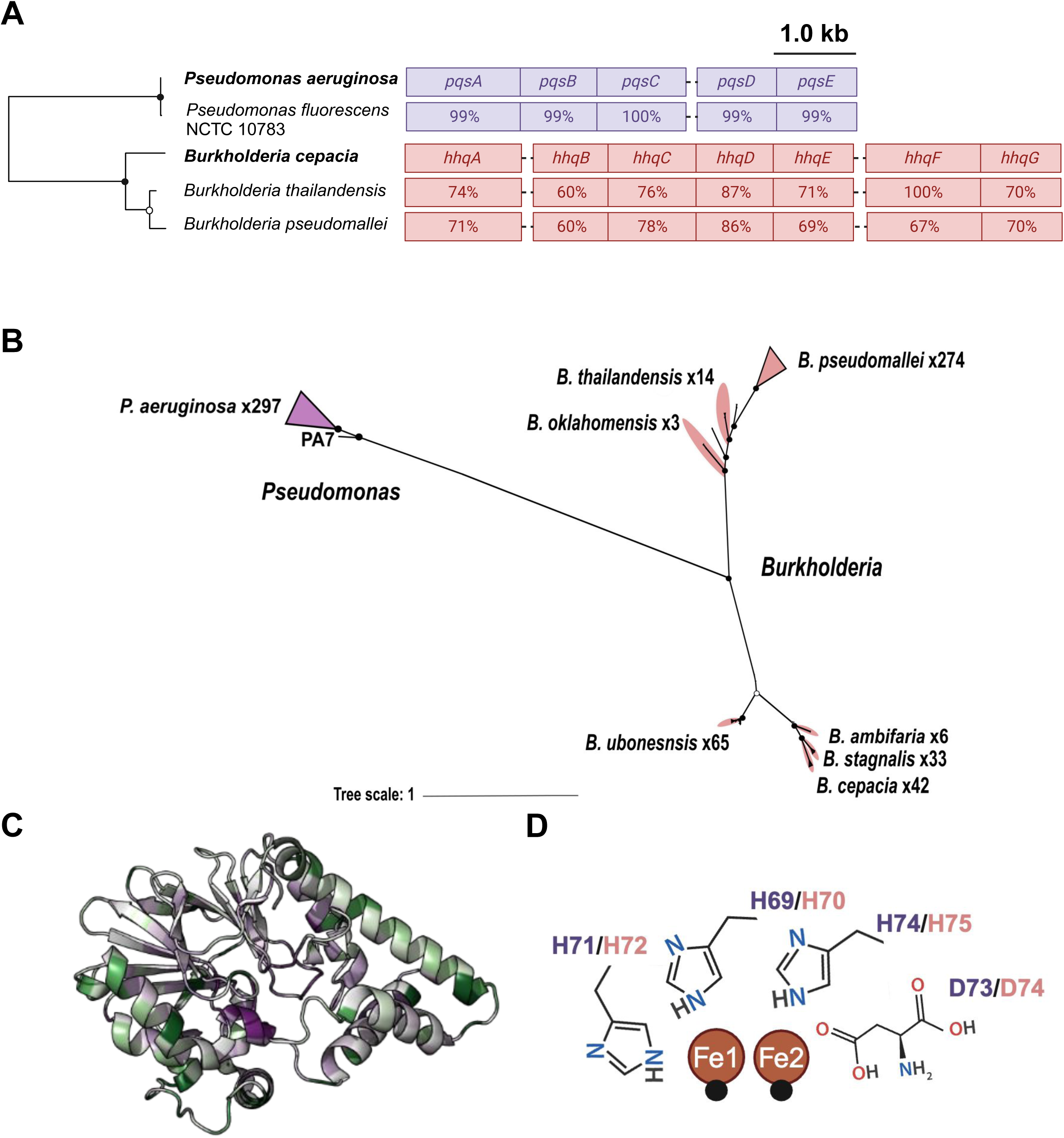
Phylogenetic analysis and conservation of PqsE and the *pqsABCDE* operon. (a) Conservation of the *pqsABCDE* and the orthologous *hhqABCDEFG* operon. Maximum-likelihood tree of PqsE/HhqE protein sequences (*n* = 7) (left). Filled and open circles represent bootstrap support values of 100% and >95%, respectively. Schematic of the *pqsABCDE* and *hhqABCDEFG* operons (right). Values represent percent identity of *pqsABCDE* from *P. fluorescens* strain NCTC 10783 compared to *P. aeruginosa* PA14 using BLASTP within the *Pseudomonas* Genome DB^33^ and *B. thailandensis* E264, *B. pseudomallei* K96243 compared to *B. cepacia* ATCC 25416. (b) Maximum-likelihood tree of orthologous PqsE protein sequences from *P. aeruginosa* and HhqE from *B. cepacia* (*n* = 764 sequences). Triangles indicate collapsed clades with 100 % sequence identity. Black circles represent bootstrap support values of 100%. A full list of sequences can be found in Data S1. (c) Cartoon representation of PqsE (PDB ID: 7KGW) colored by conservation compared with 300 homologs, estimated using the ConSurf-DB^37^. Purple residues are highly conserved and green residues are highly variable. (d) 2D projection highlighting the conserved ^69^HXHXDH^74^ motif that constitute the PqsE active site. Water molecules are depicted as black spheres, residue labels are colored dark purple based on PqsE numbering or salmon for HhqE numbering. All structure images were generated in ChimeraX^56–58^.

We identified PqsE orthologs within the *Burkholderia* genus using the *Burkholderia* Genome Database (https://www.burkholderia.com)^36^ and refer to these orthologs as HhqE. To further investigate the evolutionary relationships between these proteins, we constructed a bootstrapped phylogenetic tree that was inferred from the alignment of 761 protein sequences (Fig. 1b). This analysis revealed that PqsE proteins in *Pseudomonas* are divergent from HhqE proteins in *Burkholderia*, sharing only ∼30% amino acid identity. HhqE is highly conserved among different *Burkholderia* strains of the same species but is divergent between species. Since the function of HhqE is unknown, we performed a conservation analysis coupled with a structural analysis using the ConSurf server^37^. ConSurf predicts conserved regions based on evolutionary relationships and protein structure, allowing us to map highly conserved (functional or structurally important) residues onto the three-dimensional structure of the protein. Using an alignment of 300 PqsE orthologs, we examined which residues are conserved across *Pseudomonas* and *Burkholderia* species (Fig. 1c; Fig. S1a). The most conserved residues (purple) were located in the active site. Specifically, the ^69^HXHXDH^74^ motif responsible for the coordination of Fe^2+^/Fe^3+^ ions within the active site of PqsE was present in all orthologs (Fig. 1d). The most variable regions (green) were concentrated around the C-terminal α-helix and the loop containing residues essential for the PqsE-RhlR interaction (R170 and R171). This pattern indicates that while the PqsE-RhlR interaction may be specific to *Pseudomonas*, the catalytic core of the protein is conserved across different species, suggesting a common functional mechanism, which we test below.

### *Burkholderia* HhqE lacks the key residues for homodimerization and RhlR complex assembly

Our group and others have shown that PqsE dimerization is necessary for PqsE-RhlR complex formation^27,30^. Key residues for dimerization, located on a C-terminal α-helix unique to PqsE (compared to other metallo-β-hydrolases) have been identified in previous studies^38–39^. Notably, three arginine residues on a surface-exposed α-helix (R243/R246/R247) disrupted dimerization when substituted with alanine^27^. Importantly, this variant, which we call PqsE^NI^ for “PqsE non-interacting”, abolishes pyocyanin production, but still exhibits *in vitro* catalytic activity^27^. AlphaFold-Multimer (AFM)^40^ was used to predict structural models of HhqE proteins from *B. cepacia*, *B. pseudomallei*, and *B. thailandensis,* which aligned well with pre-existing PqsE structures, with average root mean square deviations (rmsd) of 1.80 Å, 1.90 Å, and 1.91 Å, respectively (Fig. S1b; Table S2; Movie S1). These predicted structures contain C-terminal α-helices but differ in their amino acid identities. HhqE proteins do not contain residues required for PqsE homodimerization, specifically R243, R246, R247, or E187 (Fig. 2a; Fig. S1a). Notably, in the AFM model, the residues Q187, A244, and A247, and D248 in HhqE correspond to E187, R243, R246, and R2247 in PqsE (Fig. 2b). Based on the orientation of these residues, we predicted that their absence would impact the ability of HhqE to dimerize and interact with its receptor, which we test in the following sections. HhqE proteins from *B. cepacia* (HhqE*^Bc^*) and *B. pseudomallei* (HhqE*^Bp^*) were successfully purified and selected for subsequent experimentation because of better solubility compared to other orthologs in *Burkholderia*. Size exclusion chromatogram traces established the presence of a dimeric PqsE, which eluted at an average peak fraction of 14.3 mL. In contrast, HhqE proteins eluted at later fractions: HhqE*^Bc^* eluted at an average peak fraction of 16.7 mL and HhqE*^Bp^* with an average of 17.2 mL (Fig. 2c), indicating a monomer for each of the orthologs. Complementary to size-exclusion chromatography, mass photometry (MP), a single-molecule technique, was performed to quantify the mass distribution in a given sample of PqsE and HhqE*^Bc^*. Notably, 98% of HhqE*^Bc^* counts corresponded to a monomeric form (39 kDa), supporting the chromatography results (Fig. 2d). However, PqsE exhibited dimeric counts, with 28% of counts consistent with a dimer (70 kDa) suggesting that PqsE dimerization is concentration dependent (Fig. 2d). To use the same technique but at higher concentrations, we used a microfluidic system attached to the mass photometer, which allows for the rapid dilution of a 50 µM stock protein concentration to 50 nM, allowing for the measurement of particles before they are diluted below their dissociation constant. This technique revealed a detectable population of dimers (74 kDa), representing 89% of the counts (Fig. 2e). This approach allowed us to investigate whether the oligomeric state of PqsE observed at high concentrations (50 µM) could be maintained following rapid dilution, revealing that PqsE dimerization is concentration dependent. Since PqsE dimerization is concentration dependent and dimerization is required for its ability to interact with RhlR, we hypothesize that the ability of PqsE to bind RhlR in the cell may fluctuate depending on the local PqsE concentration, potentially impacting its regulatory role in QS.

**FIG 2.**
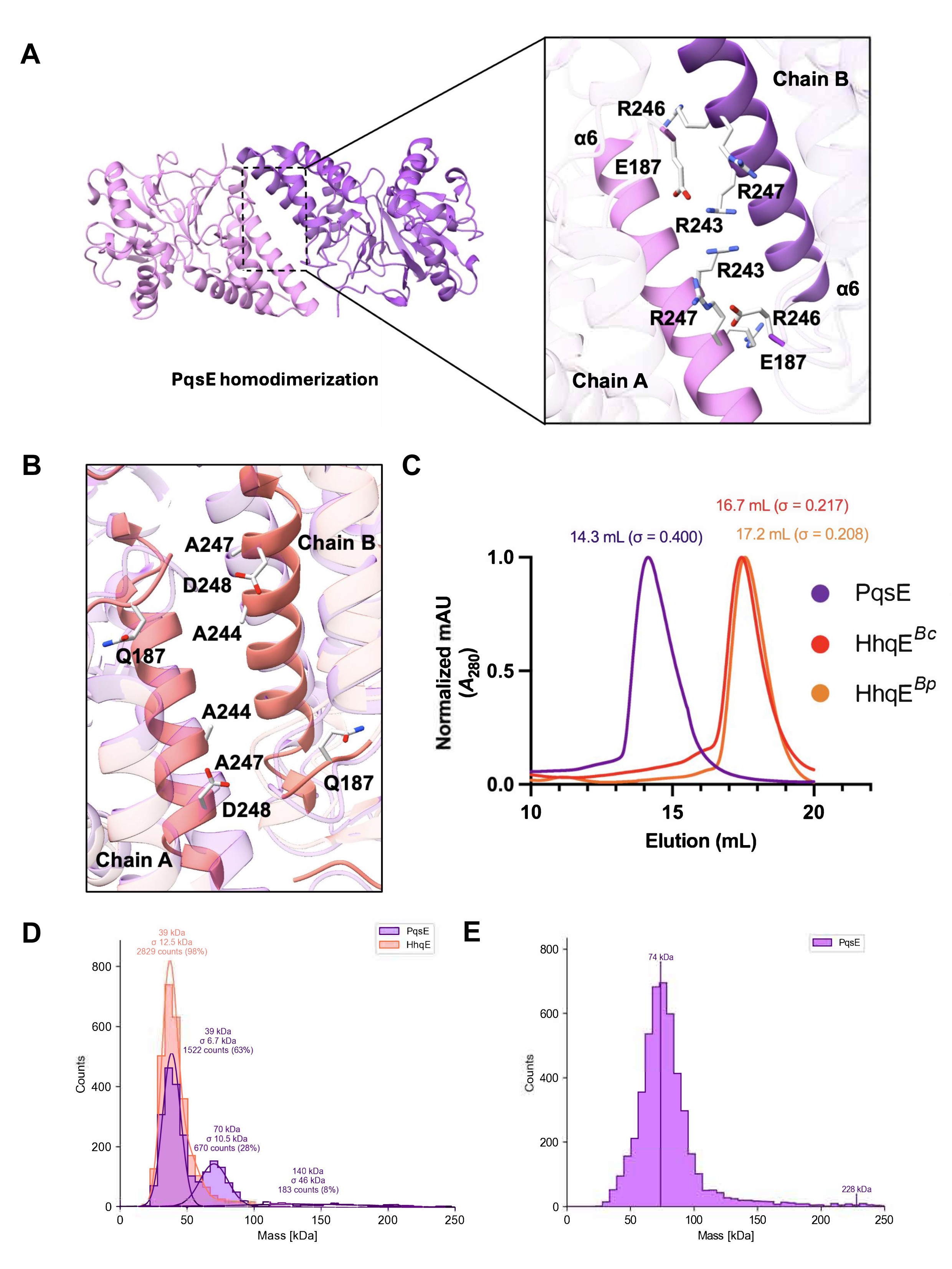
HhqE from *Burkholderia cepacia* is a monomer. (a) Cartoon representation of the PqsE homodimerization interface. Close-up view of the C-terminal interaction interface between opposing monomers. Residues E187, R243, R246, and R247 of chain A (light pink) and residues on the opposing chain (purple) are highlighted in gray and colored by heteroatom. (b) Cartoon representation of two HhqE*^Bc^*monomers (salmon) overlayed on top of the PqsE homodimer (pink). Close-up view of the HhqE*^Bc^* residues on the C-terminal α-helix that correspond to the necessary residues for PqsE dimerization. Residues Q187, A244, A247, and D248 are highlighted in gray and colored by heteroatom. (c) Size exclusion chromatography (SEC) analysis of PqsE and HhqE proteins using a Superdex-200 column, with protein elution volumes measured by absorbance at 280 nM (*A*_280_, y-axis) as a function of retention volume (mL, x-axis). Chromatogram traces are representative of six independent purifications for PqsE and HhqE*^Bc^* or three purifications for HhqE*^Bp^*. Traces are normalized to a maximum value of 1 for clarity. The mean retention volume and standard deviation (σ) are indicated above each peak. (d) Mass histograms of PqsE and HhqE*^Bc^* after manual or (e) automated dilution measured by mass photometry. Mass photometry mass distribution of 10 nM PqsE and HhqE*^Bc^* proteins diluted in PBS. 50 μM PqsE was measured with mass photometry after rapid dilution to 50 nM using the MassFluidix HC system (Refeyn). All structure images were generated in ChimeraX^56–58^.

We hypothesized that HhqE proteins would not interact with their respective LuxR-type receptors because they lack key dimerization residues and are monomeric. To test this hypothesis, we performed affinity purification pulldown analyses to assess the interaction between HhqE proteins and their “partner” receptors. Partner receptors are AHL-binding and a part of previously described LuxIR-type systems that share structural and sequence similarities to RhlR (Fig. S2)^41,42^. In this assay, PqsE or HhqE proteins were 6x-His-tagged and used as bait in affinity purification. RhlR was bound by exogenously supplied C_6_HSL, which is sufficient to activate, solubilize, and purify RhlR (manuscript in-progress). The *Burkholderia* receptors CepR and PmlR from *B. cepacia* and *B. pseudomallei*, respectively, were purified with their cognate ligand, C_8_HSL. As a positive control, we included WT PqsE and RhlR proteins, which we have previously shown to interact (Fig. 3a). Consistent with our hypothesis, we found that HhqE from *B. cepacia* and *B. pseudomallei* did not interact with RhlR or their “partner” LuxR-type receptors (Fig. 3a and Fig. 3b). In contrast, as shown in Fig. 3b, we observed an interaction between RhlR and PqsE*^Pf^*, a naturally occurring PqsE variant. Our results support a model in which *Pseudomonas* PqsE has evolved the ability to dimerize, a necessary feature for PqsE-RhlR complex formation, which makes it distinct from *Burkholderia* orthologs.

**FIG 3.**
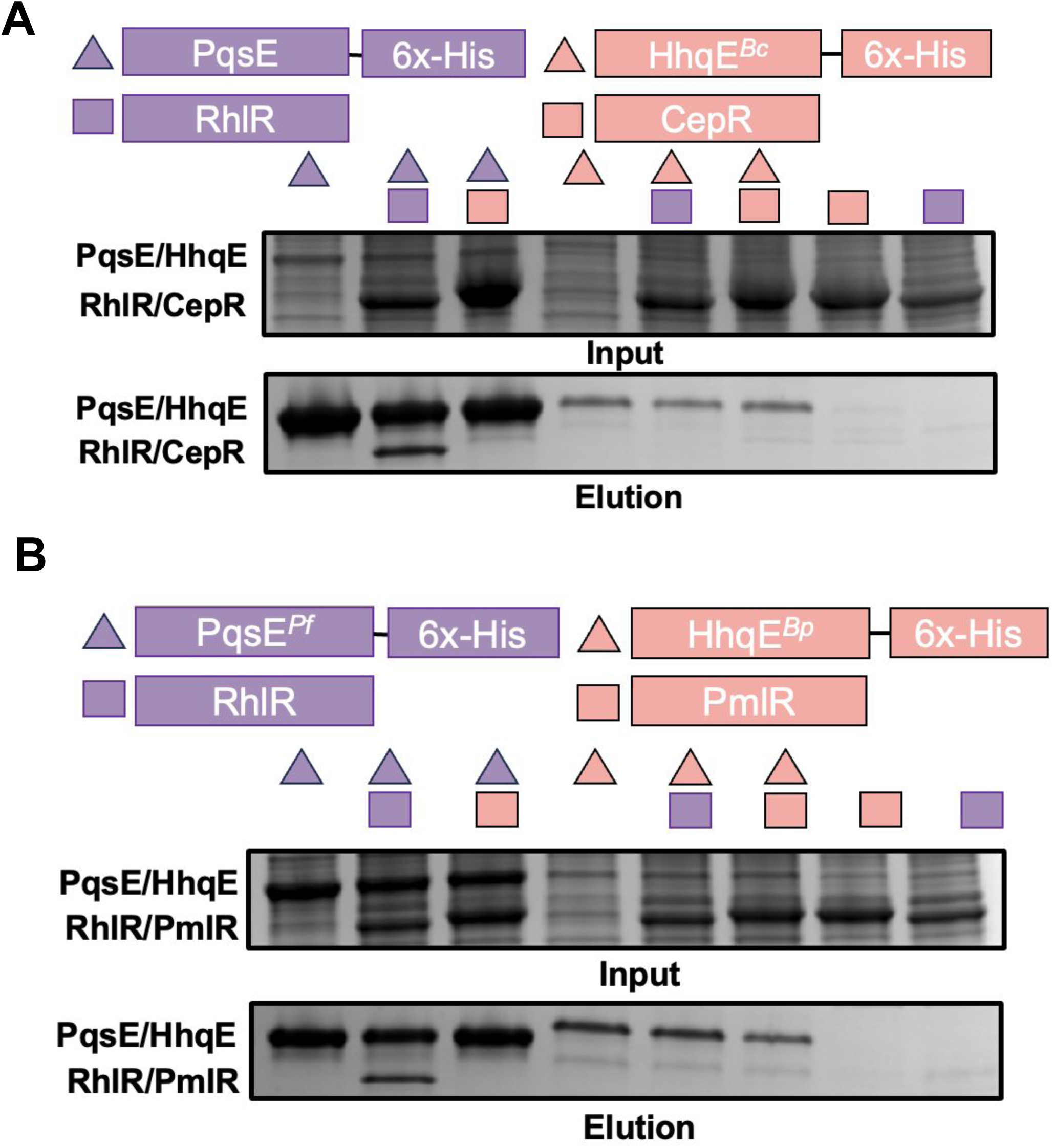
The PqsE-RhlR interaction is *Pseudomonas* specific. (a) SDS-PAGE analysis of *E. coli* cell lysates before (Input) or after (Elution) affinity purification on Ni-NTA resin. Shown are samples containing 6x-His-PqsE (purple triangles) or 6x-His-HhqE*^Bc^* (salmon triangles), either alone or in combination with *E. coli* lysates expressing RhlR:C_6_HSL (purple squares) or CepR:C_8_HSL (salmon squares). (b) Affinity purification as in (a) but using the orthologs PqsE*^Pf^* and HhqE*^Bp^*. In all affinity-purifications, RhlR and CepR lack an affinity tag and, thus, are not present in the RhlR and CepR only controls at the far right of each gel.

Consistent with not being able to interact with RhlR, we predicted that HhqE would not be able to complement the activity of PqsE in a pyocyanin production assay. We previously established that both PqsE and C_4_HSL are essential for pyocyanin production^27^; therefore, we assayed the ability of WT PqsE and HhqE*^Bc^* to produce pyocyanin in a Δ*pqsE* strain of *P. aeruginosa*. To do this, we cloned *pqsE* or *hhqE* into pUCP18 under the control of the *lac* promoter. Consistent with our current model, strains expressing *pqsE* were able to produce pyocyanin, as measured by OD_695_ and by liquid chromatography-mass spectrometry (LC-MS) (Fig. S3). In contrast, HhqE orthologs, which were unable to interact with RhlR *in vitro*, failed to complement pyocyanin production *in vivo* (Fig. S3).

### PqsE dimerization alters *in vitro* catalytic activity in a substrate-dependent manner

PqsE exhibits measurable *in vitro* enzymatic activity with synthetic substrates, enabling the evaluation of catalysis. Notably, the fluorescent substrate, 4-methylumbelliferone (MU-butyrate), is convenient because it contains a benzoic acid ring akin to the benzoate molecule that was co-crystallized in the PqsE active site and provides a direct readout for PqsE hydrolysis. Previous studies demonstrated comparable PqsE catalysis *in vitro* when comparing PqsE alone and in complex with RhlR, indicating that the PqsE-RhlR interaction does affect PqsE catalysis^27^. Moreover, catalytically inactive PqsE variants, such as PqsE D73A, retain their ability to interact with RhlR, while PqsE^NI^ maintains catalytic activity, albeit at reduced efficacy^27^. Therefore, the two primary functions of PqsE, catalysis and interaction with RhlR, can be decoupled. We used HhqE in catalysis assays to understand 1) whether HhqE*^Bc^* contains similar *in vitro* catalytic activity to PqsE and 2) whether dimerization plays a role in *in vitro* catalysis. We found that WT PqsE and HhqE*^Bc^* readily hydrolyzed MU-butyrate with different affinities and turnover rates (Table 1). While WT PqsE had a Michaelis constant (*K*_m_) of 8.1 μm, which is consistent with previous studies^27^, HhqE*^Bc^* exhibited a significantly lower *K*_m_ of 0.16 μm, indicating a higher binding affinity for MU-butyrate. However, HhqE demonstrated slower turnover. These findings suggest that while HhqE binds MU-butyrate more effectively, its catalytic efficiency is much lower compared to PqsE. We hypothesized that HhqE*^Bc^*might have a higher enzyme specificity because it is monomeric compared to PqsE. To test this, we utilized our PqsE^NI^ variant which is unable to dimerize and found comparable enzyme specificity (*K*_cat_/*K*_m_) for HhqE*^Bc^* and PqsE^NI^, suggesting that the role of the monomeric form of these proteins is enzymatic.

**Table 1.** Kinetic parameters for PqsE, HhqE and the non-interacting variant of PqsE (PqsE-NI).

| Protein | $K_m$ ($\mu\text{M}$) | $k_{\text{cat}}$ ($\text{s}^{-1}$) | $k_{\text{cat}}/K_m$ ($\text{M}^{-1} \text{s}^{-1}$) |
| --- | --- | --- | --- |
| PqsE | 8.1 | 0.0197 | $2.4 \times 10^3$ |
| HhqE | 0.16 | 0.0042 | $2.6 \times 10^4$ |
| PqsE-NI | 2.2 | 0.0562 | $2.6 \times 10^4$ |

**Acetoacetyl-CoA**
| Protein | $K_m$ (mM) | $k_{\text{cat}}$ ($\text{s}^{-1}$) | $k_{\text{cat}}/K_m$ ($\text{M}^{-1} \text{s}^{-1}$) |
| --- | --- | --- | --- |
| PqsE | 2.9 | 0.4378 | $1.5 \times 10^2$ |
| HhqE | 7.5 | 0.3115 | $4.2 \times 10^1$ |
| PqsE-NI | 2.9 | 0.3501 | $1.2 \times 10^2$ |

To further investigate the catalytic properties of PqsE and HhqE*^Bc^*, we measured the kinetic parameters for acetoacetyl-CoA hydrolysis by monitoring the release of free thiols using Ellman’s reagent. Acetoacetyl-CoA, a β-keto-thioester, was selected as a substrate because it mimics structural features of biologically relevant intermediates such as 2-ABA-CoA but lacks a benzoate ring. WT PqsE and HhqE*^Bc^* hydrolyzed this substrate with varying efficiencies (Table 1). Both enzymes exhibited weak binding affinities (high *K*_m_ values) but high turnover rates (*K*_cat_) with acetoacetyl-CoA. Our kinetic findings are consistent with previous data^26^ and demonstrate that HhqE*^Bc^*, like PqsE, prefers β-keto-thioester substrates. These results suggest that the presence of a benzoate ring, as in MU-butyrate, improves substrate binding affinity. Specifically, lower *K*_m_ values were observed for MU-butyrate compared to acetoacetyl-CoA, although efficient substrate turnover (high *K*_cat_) still required a β-keto-thioester (Table 1). This result aligns with the catalytic role of PqsE *in vivo*. Regarding the impact of dimerization on catalysis, both HhqE*^Bc^* and PqsE^NI^ displayed similarly high enzyme specificity (*K*_cat_/*K*_m_) for MU-butyrate compared to WT PqsE. However, this effect was not consistent across the tested substrates, suggesting that dimerization may influence substrate binding and turnover in a substrate-specific manner.

### RhlR ligand sensitivity reveals the need for an accessory protein

Most LuxR-type receptors fold around their cognate AHL ligands. Therefore, protein dimerization, solubility, and DNA binding occur only when ligand is present. To understand which AHL is sufficient for solubilization of our receptors of interest, RhlR and CepR, we performed protein solubility assays in the presence of cognate and non-cognate AHL (Fig. 4a). To test this, we grew *E. coli* producing either RhlR or CepR in the presence of saturating amounts of AHL, supplemented at the time of induction. Consistent with previous results^14,15^, in the absence of any ligand (DMSO only control) protein is expressed and present in the whole-cell lysate but absent in the soluble fraction. In the case of RhlR, we found that none of the AHL compounds tested were sufficient for protein solubilization when only supplied during protein induction (Fig. 4a: top gel). Soluble RhlR protein was present only when AHL was added both at the time of induction and again during cell lysis (Fig. 4a: middle gel). In contrast, medium to longer AHL chains, C_6_HSL, C_8_HSL, C_10_HSL, and 3OC_12_HSL were sufficient for solubilization of CepR. While excess amounts of cognate AHL (C_4_HSL) were not sufficient for RhlR solubilization, excess C_8_HSL, the cognate AHL for CepR, resulted in the greatest solubility for CepR.

**FIG 4.**
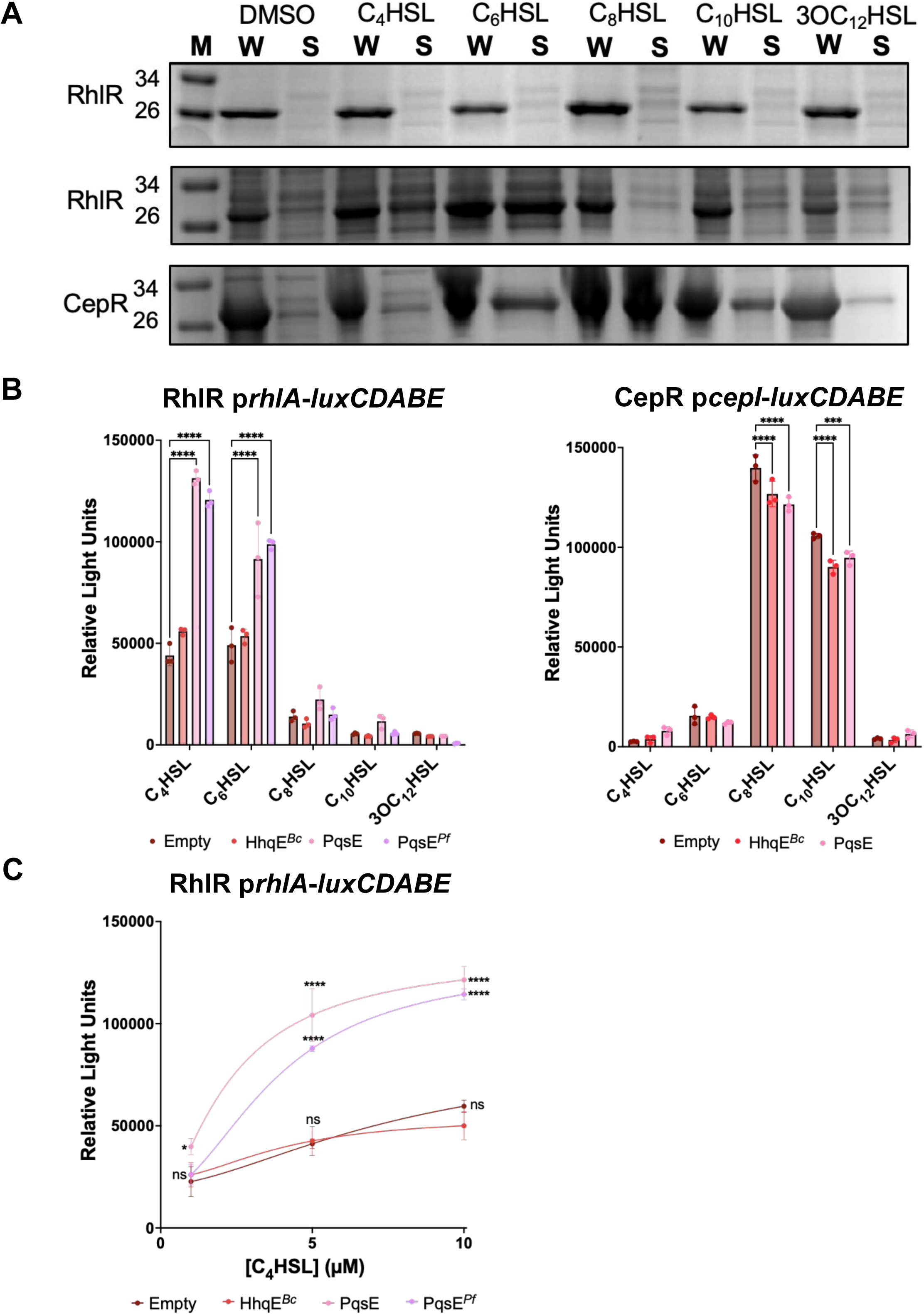
Impact of AHL and accessory protein on protein solubility and transcriptional activation in *E. coli*. (a) SDS-PAGE analysis of RhlR and CepR protein levels in whole-cell lysates (W) and soluble fractions of *E. coli* expressing each receptor from plasmids. The top and bottom gels show samples with AHL added at induction only, while the middle gel shows samples with AHL added at both induction and lysis. Protein expression was induced with 1 mM IPTG and either 1 % DMSO or 100 μM of the specified AHL. Each lane was loaded with 50 mg/mL of total protein, as measured by *A*_280_. Images are representative of three independent experiments. (b) *E. coli* reporter system expressing RhlR (left panel) or CepR (right panel) with or without accessory proteins PqsE, PqsE*^Pf^*, HhqE*^Bc^*, or empty vector control, in the presence of *rhlA* (RhlR panel) or *cepI* (CepR panel). All samples were supplemented with 0.1 % arabinose and 10 µM AHL. Bars represent the mean of three biological replicates. Error bars represent the standard deviation. Statistical analyses were performed using a two-way ANOVA with multiple comparisons using the “empty vector” as a control group within each AHL. p value summary: **** < 0.0001; non-significant comparisons are not shown. (c) Relative light unit (RLU) measurements from the RhlR reporter system co-expressed with PqsE, PqsE*^Pf^*, HhqE*^Bc^*, or empty vector control, with a three-point C_4_HSL dose response (1 µM, 5 µM, and 10 µM). Each dot represents the mean of three biological replicates. Error bars represent the standard deviation. Statistical analyses were performed using a two-way ANOVA with multiple comparisons using the “empty vector” as a control group for each concentration. p value summary: * < 0.05; *** < 0.001; **** < 0.0001; ns = not significant.

The results in Fig. 4a suggest that RhlR has a narrow range of AHL that can activate RhlR transcription. In contrast, we expected that CepR will have a broader range of AHL-dependent transcriptional activation, based on the ability of cognate and non-cognate AHL to solubilize CepR. To test this, we employed a three plasmid *E. coli* reporter system. This recombinant *E. coli* strain harbors one plasmid with arabinose-inducible expression of *rhlR* or *cepR,* and a second plasmid with the luciferase operon fused to a promoter controlled by the regulator of interest, p*rhlA* and p*cepI*, respectively. This reporter was previously used to demonstrate that PqsE, constitutively expressed on a third plasmid, enhances RhlR-dependent transcription in the presence of C_4_HSL^27–29^. However, it remains unclear if PqsE has a similar effect across various AHL-chains or if this stimulatory effect can be complemented by PqsE*^Pf^* or *Burkholderia* HhqE*^Bc^*. Consistent with previous findings, we observed enhanced RhlR-dependent transcription upon the addition of increasing concentrations of C_4_HSL, which was enhanced in the presence of PqsE and PqsE*^Pf^* (Figs. 4b; left panel)^27–29^. Additionally, C_6_HSL activated RhlR-dependent transcription, with further enhancement observed in the presence of PqsE. Conversely, co-expression of HhqE with RhlR mirrored the transcriptional response observed in the presence of an empty vector control, indicating that HhqE does not enhance RhlR-dependent transcription. CepR responded to its cognate ligand C_8_HSL and, to a lesser extent, C_10_HSL (Fig. 4b; right panel). These results are consistent with protein solubility assays where medium length AHL-chains were sufficient for CepR folding. The presence of PqsE or HhqE did not further increase CepR-dependent transcription, consistent with our *in vitro* pulldown assays. We take this to mean that AHL binding to CepR is sufficient for maximum transcription and does not require an accessory protein, like RhlR with PqsE. Given that PqsE has the ability to enhance the affinity of RhlR for promoter DNA across different concentrations of AHL (Fig. 4c), we aimed to confirm that HhqE does not alter the binding affinity of CepR for its target promoter. To test this, we performed electrophoretic mobility shift assays (EMSA) to assess the binding affinity of CepR to DNA compared to CepR in the presence of HhqE. Using the CepR-dependent *cepI* promoter that was identical to the fragment used in the *E. coli* luciferase reporter assays, we found that CepR bound the promoter DNA and that the presence of HhqE did not alter this binding (Fig. S4). As a negative control, we performed the same assay using CepR with the RhlR-dependent *rhlA* promoter and observed no shift in the DNA, confirming the specificity of CepR DNA binding (Fig. S4).

### PqsE prevents degradation of RhlR by Lon protease in the absence of AHL

RhlR is relatively insoluble in the presence of AHL at the time of expression, requiring excess AHL in the lysis buffer to promote protein solubility (Fig. 4a). We hypothesized that this is because our *E. coli* solubility assays do not express PqsE to help stabilize RhlR and that it is instead rapidly turned over. To test this supposition in *P. aeruginosa*, we performed western blots against RhlR using a polyclonal antibody (raised against purified RhlR) in different genetic backgrounds to determine the role of PqsE in protecting RhlR from degradation *in vivo.* Consistent with previous studies, RhlR levels were comparable between WT and Δ*rhlI* strains, indicating that C_4_HSL alone does not contribute to RhlR stability^43^, likely because of the presence of PqsE in both of these strains (Fig. 5a). We found significantly reduced RhlR levels in Δ*pqsE* or *pqsE*^NI^ strains (Figure 5a). There was a notable reduction in RhlR levels in the Δ*rhlI* Δ*pqsE* double deletion strain. Additionally, the *pqsE*^NI^ mutation in a Δ*rhlI* background had reduced RhlR levels, suggesting that the PqsE-RhlR interaction is necessary to prevent RhlR turnover independent of C_4_HSL. Previous work in the laboratory strain PAO1 showed increased RhlR levels in cells with a disrupted ATP-dependent Lon protease^44^. We hypothesized that PqsE binding to RhlR prevents recognition of RhlR by Lon; thus, we expected that a Δ*lon* Δ*pqsE* strain would exhibit levels of RhlR comparable to WT. To test this, we constructed a *P. aeruginosa* strain with a deletion in the gene encoding the Lon protease in both the WT and Δ*pqsE* backgrounds. We found significantly increased RhlR levels in a Δ*lon* deletion strain (Fig. 5b). Additionally, RhlR levels were rescued back to WT levels in a Δ*lon* Δ*pqsE* strain. Thus, we conclude that PqsE protects RhlR from degradation by the Lon protease and the absence of PqsE leads to a decrease in RhlR levels (Fig. 5b).

**FIG 5.**
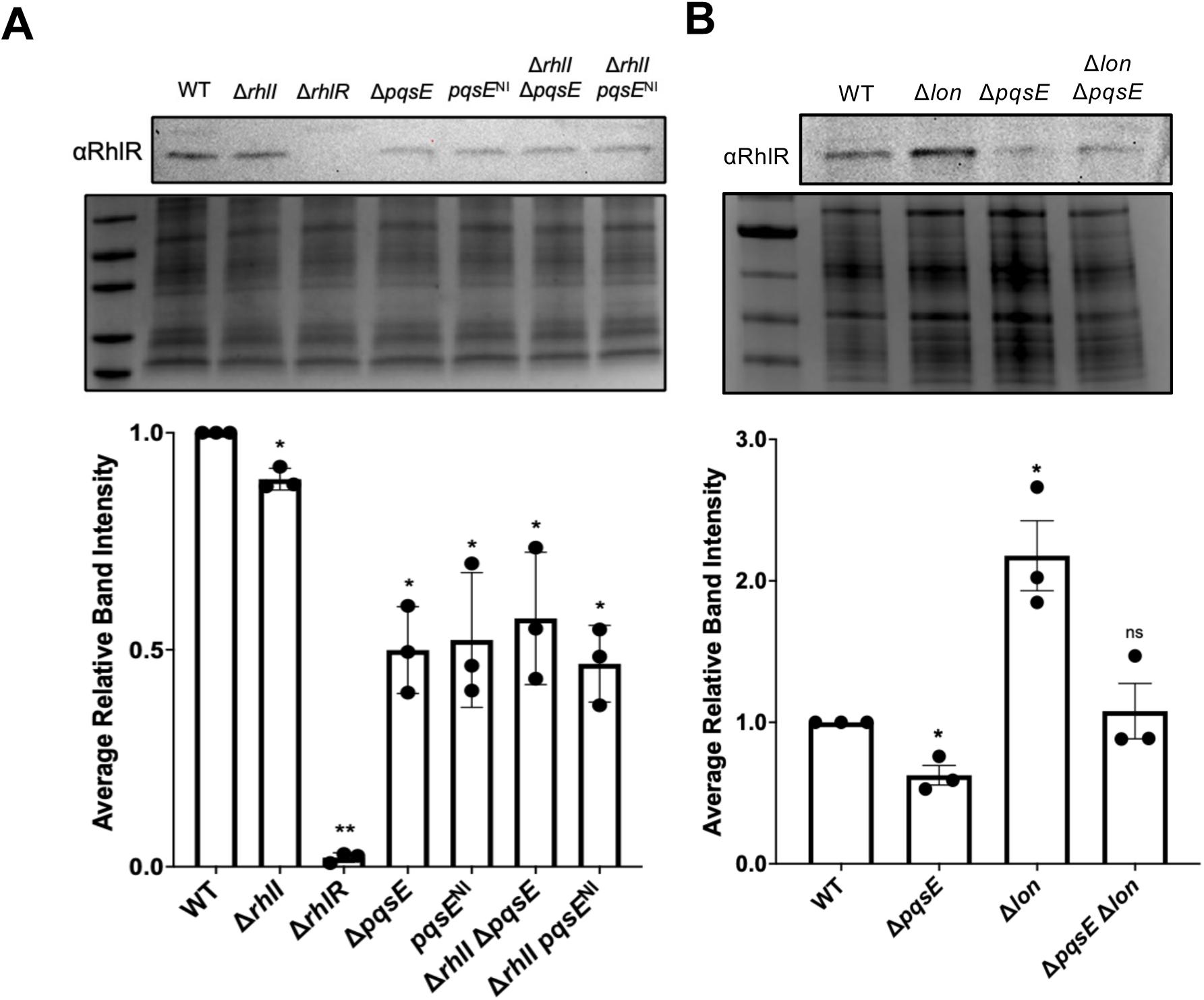
PqsE prevents targeted degradation of RhlR by the Lon protease. (a) Western blot analysis of lysates from WT PA14, Δ*rhlI*, Δ*rhlR*, Δ*pqsE*, *pqsE*^NI^, Δ*rhlI* Δ*pqsE*, and Δ*rhlI pqsE*^NI^ backgrounds. RhlR levels were detected using a polyclonal antibody against RhlR and compared to an SDS-PAGE loading control of total protein levels (bottom gel). Average relative intensities of RhlR levels in different PA14 backgrounds. (b) Same as (a) except RhlR levels were assessed in WT PA14, Δ*pqsE*, Δ*lon*, and Δ*pqsE* Δ*lon* strains. RhlR levels were determined by densitometric analysis where RhlR levels for each strain were quantified relative to its respective total protein loading control and then compared to WT PA14 RhlR levels. The data are the mean of three independent quantifications of three biological replicates. Error bars correspond to standard deviation. Statistical analyses were performed using a ratio paired t test between WT and the individual mutant strains. p value summary: * < 0.05; ** < 0.01; ns = not significant.

### Convergent evolution in *A. baumannii*: the AbaR LuxR-type receptor interacts with BlsA

Recently, our model of an accessory protein activating transcription of a LuxR-type receptor was supported by a report in *A. baumannii*^45^. In *A. baumannii*, the light-sensing protein BlsA interacts with the LuxR-type receptor AbaR, inducing the expression of its partner synthase *abaI*. To better understand this interaction and facilitate a comparison with the previously solved PqsE-RhlR structure, we generated a structural model using AFM and used this predicted structure as a guide to identify residues crucial for the BlsA-AbaR protein-protein interaction (Fig. 6a). We confirmed this interaction using a bacterial adenylate cyclase-based two-hybrid system (BACTH), which was chosen to circumvent the difficulty in expressing a soluble form of AbaR. In this assay, we used an *E. coli* strain lacking the *cyaA* gene, which encodes adenylate cyclase.

**FIG 6.**
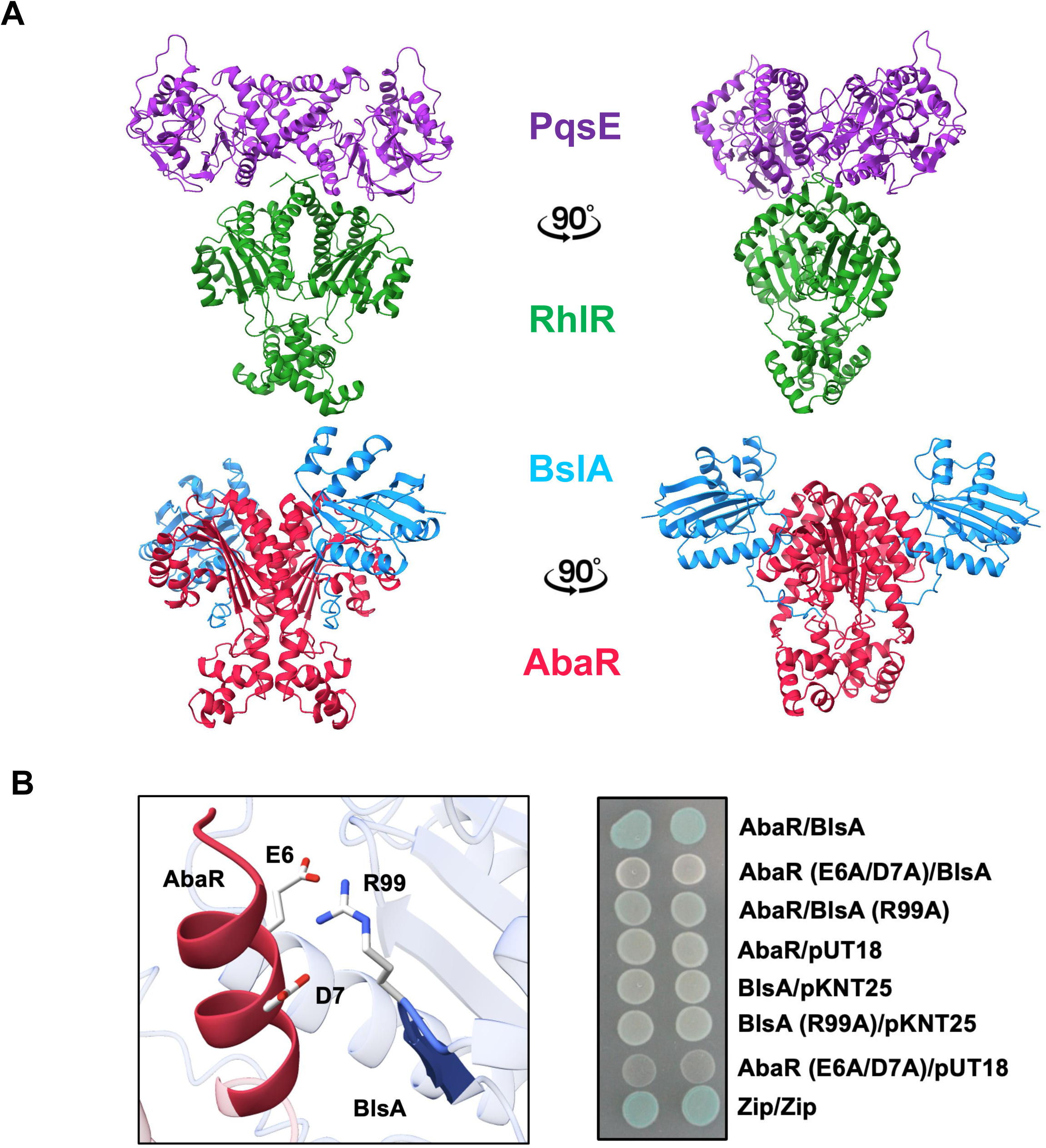
Characterization of the AbaR-BlsA interaction from *Acinetobacter baumannii.* (a) Structural comparison of the cryo-electron microscopy structure of RhlR (green) bound to PqsE (purple) (PDB ID: 8DQ0^28^) with the AlphaFold-Multimer predicted structure of AbaR (red) bound to BslA (blue). (b) Close-up view of the AbaR-BlsA predicted interaction interface, highlighting residues E6 and D7 on AbaR and residue R99 in BlsA, colored by heteroatom. Interaction between AbaR-BlsA was evaluated by using a bacterial adenylate cyclase two-hybrid (BACTH) analysis. *E. coli* MG1655 Δ*cyaA* strains harboring plasmids encoding the T25 and T18 fragments of *Bordetella pertussis* adenylyl cyclase, fused to either WT *abaR* and *blsA* or the indicated point-mutants, were assayed for interaction. All structure images were generated in ChimeraX^56–58^.

The WT and point mutants of *abaR* and *blsA* were cloned into the vectors pKNT25 and pUT18, which encode the T25 and T18 fragments of *Bordetella pertussis* adenylate cyclase, respectively. When co-expressed in the Δ*cyaA* strain of *E. coli*, physical interactions between the proteins bring the T25 and T18 domains into close proximity, reconstituting a functional adenylate cyclase activity. Interaction was detected by plating on X-Gal-containing medium, where blue colonies indicated successful cAMP production and *lacZ* expression. White colonies indicate no interaction between the proteins. As expected, WT AbaR and BlsA co-expression resulted in blue colonies, indicating an interaction between the two proteins. Little to no blue coloration was observed when either protein was co-expressed with their respective empty vector controls (Fig. 6b). Utilizing our structural model of BlsA-AbaR, we identified AbaR residues E6 and D7 as important for an electrostatic interaction with residue R99 on BlsA and generated alanine-substitution variants. Co-expression of the AbaR variant E6A/D7A with WT BlsA, as well as BlsA R99A with WT AbaR disrupted the protein-protein interaction, as indicated by the loss of blue pigment compared to WT and the positive control (Zip/Zip) (Fig. 6b). Thus, AbaR requires an accessory protein to stabilize and activate transcription, like RhlR, suggesting that AHL binding alone may not be sufficient for proper function. This dependency on an accessory protein could be related to structural features within the ligand-binding pocket (LBP), such as accessibility of the binding site to AHL. While the exact structural basis for this requirement remains unclear, we hypothesized that the size and solvent accessibility of LBP may contribute to the need for accessory factors. To explore this further, we compared well-characterized LuxR-type receptors and their solvent-accessible binding site volumes (Table 2). We found that AbaR and RhlR exhibit the smallest solvent-accessible volume, nearly half that of the next smallest receptor. This analysis provides insight into how structural differences across receptors may influence ligand binding and how dependency on accessory proteins determines different mechanisms of activation.

**Table 2.**
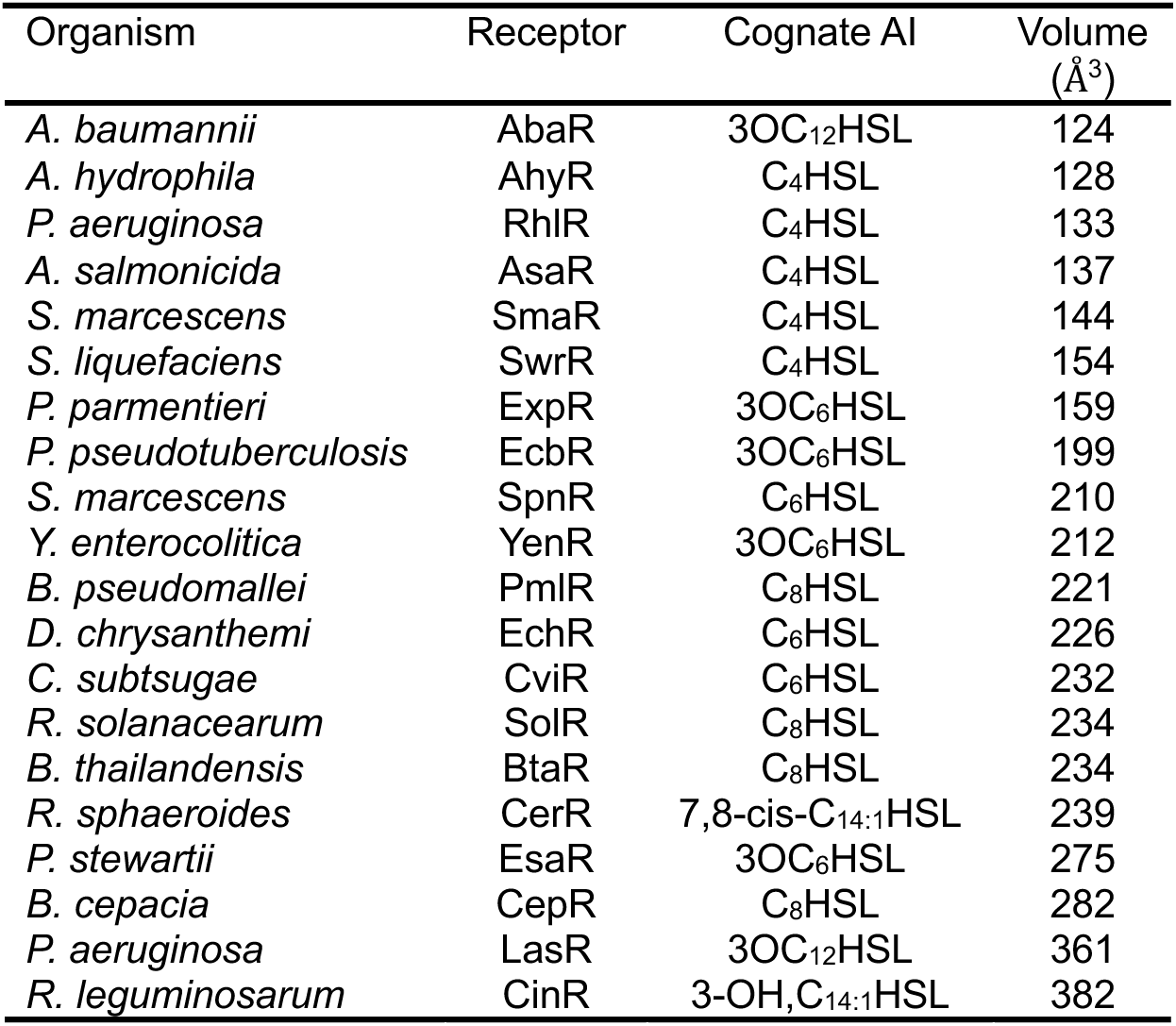
Solvent accessible volume of well-characterized LuxR-type receptors. This table includes the organism, receptor name, cognate AHL, and solvent accessible volume calculated using DoGSite3^59^ within the Proteins*Plus* web server^60^.

| Organism | Receptor | Cognate AI | Volume<br>(Å<sup>3</sup>) |
| --- | --- | --- | --- |
| *A. baumannii* | AbaR | 3OC<sub>12</sub>HSL | 124 |
| *A. hydrophila* | AhyR | C<sub>4</sub>HSL | 128 |
| *P. aeruginosa* | RhlR | C<sub>4</sub>HSL | 133 |
| *A. salmonicida* | AsaR | C<sub>4</sub>HSL | 137 |
| *S. marcescens* | SmaR | C<sub>4</sub>HSL | 144 |
| *S. liquefaciens* | SwrR | C<sub>4</sub>HSL | 154 |
| *P. parmentieri* | ExpR | 3OC<sub>6</sub>HSL | 159 |
| *P. pseudotuberculosis* | EcbR | 3OC<sub>6</sub>HSL | 199 |
| *S. marcescens* | SpnR | C<sub>6</sub>HSL | 210 |
| *Y. enterocolitica* | YenR | 3OC<sub>6</sub>HSL | 212 |
| *B. pseudomallei* | PmlR | C<sub>8</sub>HSL | 221 |
| *D. chrysanthemi* | EchR | C<sub>6</sub>HSL | 226 |
| *C. subtsugae* | CviR | C<sub>6</sub>HSL | 232 |
| *R. solanacearum* | SolR | C<sub>8</sub>HSL | 234 |
| *B. thailandensis* | BtaR | C<sub>8</sub>HSL | 234 |
| *R. sphaeroides* | CerR | 7,8-cis-C<sub>14:1</sub>HSL | 239 |
| *P. stewartii* | EsaR | 3OC<sub>6</sub>HSL | 275 |
| *B. cepacia* | CepR | C<sub>8</sub>HSL | 282 |
| *P. aeruginosa* | LasR | 3OC<sub>12</sub>HSL | 361 |
| *R. leguminosarum* | CinR | 3-OH,C<sub>14:1</sub>HSL | 382 |

## DISCUSSION

Our study revealed key functional differences between PqsE from *P. aeruginosa* and HhqE orthologs from *Burkholderia* species, primarily due to the unique ability of PqsE to dimerize. Previous research mapped the distribution of the *hhqABCDEFG* operon, which includes HhqE orthologs across the *Burkholderia* genus^21,46^ However, little was known about the specific functional roles of these orthologs compared to PqsE. Interestingly, orthologs of PqsE were found exclusively in *Pseudomonas* and *Burkholderia*, suggesting evolutionary divergence. Although previous studies have not found direct sequence-level evidence of horizontal gene transfer (HGT), we propose that it is more likely that *Pseudomonas* acquired PqsE through HGT than the intermediate lineages to have lost it. We demonstrated that while PqsE and HhqE share structural similarities, only PqsE exhibits concentration-dependent dimerization, which is critical for its interaction with the LuxR-type receptor RhlR and the activation of QS virulence factors. Our results suggest that dimerization is key to the regulatory role of PqsE in RhlR-dependent transcription. Size-exclusion chromatography and MP revealed that PqsE is monomeric at low concentrations, but forms dimers at higher concentrations. This equilibrium suggests that PqsE-RhlR complex formation is dependent on PqsE levels, allowing *P. aeruginosa* to regulate virulence in response to environmental or QS signals. In contrast, HhqE proteins from *Burkholderia* species, lack the key residues necessary for dimerization (R243, R246, and R247), and remain monomeric at all tested concentrations. This inability to dimerize may explain the failure of HhqE proteins to interact with its respective LuxR-type receptor, as confirmed by pull-down assays. Concentration-dependent dimerization of PqsE suggests a model in which its activity is modulated by environmental conditions. At low concentrations, PqsE may primarily function as a monomer with enzymatic activity independent of RhlR. However, at higher concentrations, such as during the late stages of infection, PqsE dimerization can promote its interaction with RhlR, enhancing the expression of virulence factors such as pyocyanin. The divergence of HhqE, which remains monomeric and is incapable of LuxR-type receptor interaction, underscores its functional specialization in *Burkholderia*. While HhqE retains its catalytic function *in vitro*, the lack of transcriptional regulation via receptor interactions suggests that it plays a more limited role in QS than PqsE. This specialization may reflect distinct evolutionary paths for QS and virulence regulation in *Pseudomonas* and *Burkholderia*. Previous studies have shown that Δ*pqsE* strains of *P. aeruginosa* produce WT levels of PQS, likely due to other thioesterases, such as TesB, that compensate for the loss of PqsE in PQS biosynthesis^26^. This suggests that the primary role of PqsE is its interaction with RhlR, rather than its thioesterase activity. In contrast, *Burkholderia* species lack a TesB homolog, further supporting the idea that HhqE primarily functions in catalysis, without forming complexes with LuxR-type receptors. This functional divergence suggests that *Burkholderia* spp. may rely on alternative mechanisms for QS and virulence regulation, distinct from the transcriptional activation observed in *Pseudomonas*.

We propose that the solvent-accessible area of the ligand-binding pocket (LBP) plays a critical role in determining ligand specificity. A larger solvent-exposed LBP can accommodate a broader range of AHL molecules, facilitating more promiscuous binding, while a more enclosed LBP restricts ligand access and enforces specificity. Supporting this hypothesis, the LBPs of AbaR, RhlR, and LasR exhibit distinct differences in size, hydrophobicity, and hydrogen-bonding properties that may influence ligand specificity and receptor function. LasR, with the largest and most solvent-accessible LBP, has a hydrophobicity ratio of 0.752, a balanced number of H-bond donors and acceptors (12 donors, 15 acceptors), and a hydrophobic composition with 30 aromatic atoms, four alanine and five leucine residues, supporting its ligand promiscuity. In contrast, AbaR and RhlR have restricted LBPs, with higher hydrophobicity ratios (0.793 and 0.777, respectively), fewer aromatic atoms (27 and 33), and reduced H-bond donors and acceptors, likely contributing to their limited ability to accommodate diverse AHLs. Interestingly, both AbaR and RhlR require accessory proteins for transcriptional activation, suggesting a potential evolutionary trade-off between ligand specificity and receptor stability and function. Their constrained LBPs and reduced hydrophobic residues may limit flexibility and necessitate accessory protein-mediated function. These observations imply that variations in the solvent-accessible area of the LBP have evolved to balance detection of diverse signals, as observed in LasR^15^ and CepR (Fig. 4), with regulatory precision and control, as seen in AbaR and RhlR. Future studies investigating LBP architecture and accessory protein interactions will be crucial in understanding the mechanistic diversity of LuxR-type transcription factors.

The *P. aeruginosa* hierarchical quorum-sensing network highlights distinct roles for LasR and RhlR in regulating virulence and gene expression. LasR, with its larger, more promiscuous ligand-binding pocket, responds to a wide range of signals early in quorum sensing, initiating virulence gene expression. In contrast, RhlR, with a more restricted and specific ligand-binding pocket, functions downstream, fine-tuning gene expression at later stages. RhlR activity is further regulated by PqsE, which enhances RhlR-dependent gene expression, introducing an additional level of control. We hypothesize that this hierarchical arrangement allows LasR to act as a generalist sensor for broad environmental cues, while RhlR serves as a specialist to refine responses, ensuring precise timing and coordination of virulence factor production as quorum sensing progresses. More specifically, we postulate that there is a narrow window that promotes RhlR activation. Approximately 1-5 μM of C_4_HSL is required to activate RhlR^47^, which is an order of magnitude higher than the levels required by other QS receptors^15,48^, such as LasR with 3OC_12_HSL. It is possible this high threshold for activation is a mechanism to ensure that RhlR-dependent traits are expressed only at sufficient cell density. Consequently, without sufficient levels of AHL to stabilize the receptor, RhlR would be vulnerable to turnover by the proteasome. However, *P. aeruginosa* has specifically evolved PqsE to provide a mechanism to protect RhlR from degradation and to selectively drive it to promoters to drive differential gene expression^29^.

Our data suggest a key role for PqsE in protecting RhlR from degradation by the Lon protease, which influences QS progression in *P. aeruginosa*. Western blot analyses showed that RhlR levels significantly decreased in strains lacking PqsE or in strains where PqsE is unable to dimerize (Fig. 5a). This reduction was reversed in a Δ*lon* Δ*pqsE* strain, indicating that, in the absence of PqsE, RhlR becomes a target of Lon protease-mediated degradation (Fig. 5b). In our proposed model, at high cell density (HCD), RhlR is bound by PqsE, enhancing the production of virulence factors, like pyocyanin in response to elevated C_4_HSL levels. This interaction between PqsE-RhlR not only promotes QS-regulated gene expression but also shields RhlR from degradation by Lon protease, stabilizing RhlR during active QS (Fig. 7). As cell density increases, we hypothesize that waning levels of PqsE lead to fewer PqsE-RhlR complexes in the cell, allowing Lon protease to target free RhlR. Since PqsE dimerization is concentration dependent (Fig. 2d), its ability to bind RhlR may fluctuate in response to cellular concentrations of PqsE, potentially influencing the stability of RhlR. This concentration dependence fits into our model, suggesting that as PqsE levels decline, dimerization becomes insufficient to prevent RhlR from Lon-mediated degradation. This Lon-mediated degradation of RhlR may play a crucial role in transitioning from HCD-associated group behaviors to low-cell density (LCD) states, facilitating reprogramming of cellular activities as AHL signals dissipate. While little is known about the mechanisms underlying the HCD-to-LCD transition in *P. aeruginosa*, studies in *Vibrio cholerae* have provided key insights into this process. In *V. cholerae*, the transition from HCD to LCD is accompanied by biofilm dispersal and repression of QS-regulated behaviors^49^. Additionally, the transcriptional regulator SmcR in *V. vulnificus* is degraded by the ClpAP protease, setting precedence for targeted degradation of these regulators in QS systems^50^. We propose that the Lon protease may play a role in *P. aeruginosa* by selectively degrading RhlR once PqsE levels decline, facilitating a transition away from RhlR-dependent behaviors at HCD. Investigating how Lon protease modulates this transition and identifying the specific conditions under which it targets RhlR could provide insights into the regulatory mechanisms underlying QS downregulation. Further exploration could also identify potential regulatory checkpoints within QS that impact the pathogenicity and virulence of *P. aeruginosa*.

**FIG 7.**
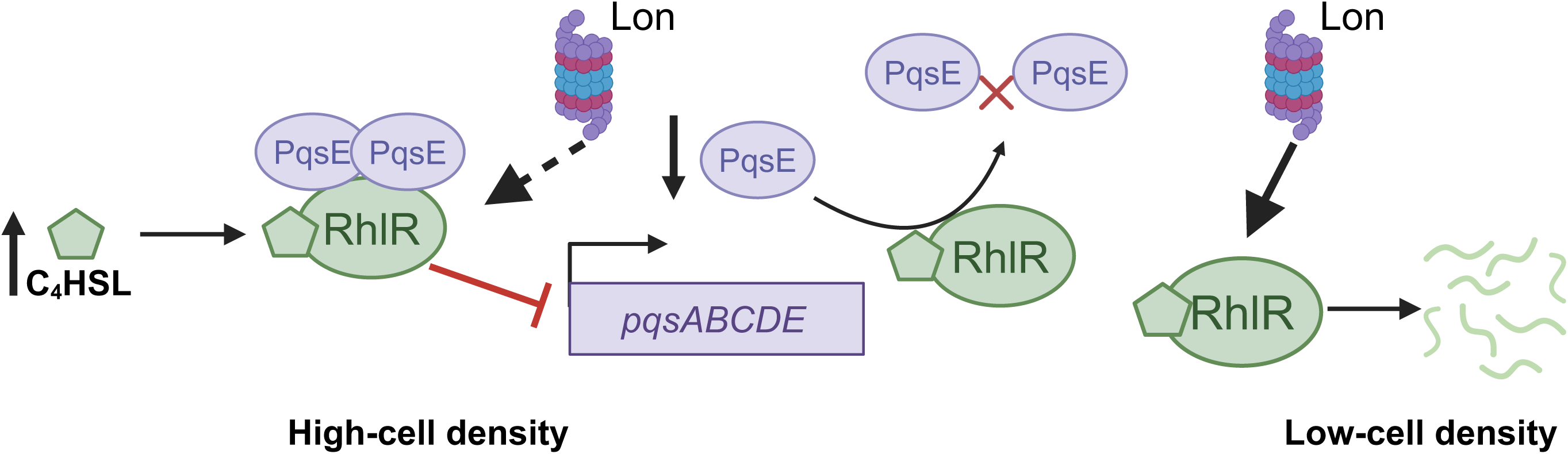
Model for targeted degradation of RhlR by the Lon protease. RhlR bound by PqsE is not targeted by degradation. At high cell density, C_4_HSL levels increase leading to increased activation of RhlR and repression of the *pqs*ABCDE operon. The repression of *pqsE* leaves RhlR unprotected and more prone to degradation by Lon protease, thus reverting signaling back to low-cell density behaviors. Schematic was made using BioRender.

## MATERIALS AND METHODS

### Strain construction

Genes encoding *hhqE* and *cepR* were amplified from *B. cepacia* (ATCC 25416) genomic DNA using traditional PCR-based methods. Orthologs from *P. fluorescens* strain NCTC 10783, *B. thailandensis* E264, and *B. pseudomallei* strain K96243 were cloned from gene fragments obtained from Twist Bioscience. Cloning for these constructs was conducted following a similar protocol. Cloning for some constructs were carried out using Hifi DNA assembly (NEBuilder) where appropriate. To construct a marker-less in-frame chromosomal deletion in PA14, two 500 bp regions flanking a 21 bp segment of the *lon* coding sequence were amplified from a gene fragment (Twist Bioscience). This fragment was cloned into the pEXG2 vector and subsequently transformed into a conjugation-competent *E. coli* SM10 λ*pir* strain. Conjugation was carried out with WT PA14 and Δ*pqsE* strains of *P. aeruginosa*. A detailed list of strains, plasmids, and oligonucleotide primers used in this study is provided in Table S2.

### Protein production and purification

The expression of PqsE variants and orthologs was performed in *E. coli* BL21 (DE3) cells transformed with the gene of interest cloned into the pET28b vector. Overnight cultures from glycerol stocks were diluted 1:100 in LB medium supplemented with kanamycin (50 μg/mL) and grown to OD_600_ of 1.0 at 37 °C with shaking at 200 rpm. Protein expression was induced with 1 mM isopropylβ-D-1-thiogalactopyranoside (IPTG). For PqsE variants, cultures were grown for an additional 4 h at 25 °C with 200 rpm. HhqE proteins were induced at 18 °C and grown overnight with shaking at 130 rpm. We observe that for every 1 L of PqsE culture, 2 L of HhqE was required to achieve comparable yields of soluble protein, as lower volumes yielded insufficient amounts of HhqE. CepR expression followed a similar protocol. Full-length CepR was cloned into the pET-DUET vector and overexpressed in *E. coli* BL21 cells. Cultures were grown at 30 °C until reaching an OD_600_ of 0.6, at which point protein production was induced with 0.5 mM IPTG in the presence of 100 μm C_8_HSL. CepR cultures were grown at 18 °C overnight with shaking 130 rpm.

### Phylogenetic analyses

To investigate the evolutionary relationship between PqsE and HhqE protein groups, we first identified orthologs using the *Pseudomonas* Genome DB^33^ and *Burkholderia* Genome DB^36^ respectively. The two lists were concatenated and then aligned using MUSCLE v3.8.1551^51^ and trimmed with trimAl v1.4.1^52^. The maximum-likelihood tree was generated from an alignment of (*n* = 762) PqsE and HhqE sequences with IQ-TREE v1.6.12^53^. Model selection was performed using an automatic substitution model based on the Bayesian information criteria (BIC) score, where the WAG+F+G4 model was chosen, and bootstrap support values were performed with 1,000 replicates. The tree was visualized and annotated with Interactive Tree of Life (iTOL) v6.9.1^54^.

### Pyocyanin assays

*P. aeruginosa* Δ*pqsE* strains carrying either the pUCP18 vector or the vector containing WT *pqsE* or *hhqE* genes, were cultured from glycerol stocks in 5 mL of LB + carbenicillin 400 μg/mL at 37°C with shaking. Overnight cultures were back diluted 1:10 in 1 mL of LB, and cell density was measured at OD_600_. The remaining 1 mL of each culture was centrifuged at (21,300 x *g*) for 2 min, and the absorbance of cell-free supernatants was measured at OD_695_ to quantify pyocyanin. Cell-free supernatants were frozen for ultra-high-performance liquid chromatography coupled (UHPLC) with high-resolution mass spectrometry and analyzed as described previously^47^.

### Mass photometry

Mass photometry experiments were carried out using a TwoMP system (Refeyn Ltd.). Samples were initially diluted in PBS, then further diluted 1:10 into a 20 μl droplet placed in the well of a silicon gasket attached to the instrument, using a glass coverslip coated with poly-L-lysine. The final sample concentration was 10 nM. For the mass fluidics experiment, PqsE was prepared at an initial concentration of 50 μM in PBS and analyzed using the MassFluidix HC system (Refeyn Ltd.). PqsE was subject to rapid dilution with a dilution factor of 1:1000, achieving measurement on the millisecond timescale. For contrast to mass conversion, bovine serum albumin (66-kDa) and immunoglobulin G (150- and 300-kDa) were used as calibrants on the same day as each measurement. Data were analyzed using DiscoveryMP (Refeyn Ltd.).

### Affinity-purification pulldown

*E. coli* strains harboring overexpression vectors producing 6x-His PqsE orthologs or un-tagged RhlR orthologs were grown overnight and back-diluted 1:100 in LB + ampicillin (100 μg/mL) or kanamycin 50 μg/mL). Cultures were grown to OD_600_ of 1.0 at 37 °C with 200 rpm. Protein production was induced with 1 mM IPTG. In the case of RhlR, CepR, and PmlR, 150 μm of C_6_HSL or C_8_HSL were added at the time of induction. Induced cultures were incubated at 18 °C overnight with shaking at 130 rpm and cells were harvested the following day at 10,876 x *g*. Cell pellets were resuspended in 1 mL of lysis buffer (20 mM Tris-HCl pH 8.0, 150 mM NaCl) and lysed by sonication (3 x 30-s pulses at 25% amplitude). In the case of RhlR, C_6_HSL was supplemented to the lysis buffer at a final concentration of 150 μm to ensure soluble protein. Following sonication, lysed cells were transferred to microcentrifuge tubes and subjected to centrifugation at 21,300 x *g* at 4 °C for 20 min. Supernatant fractions containing PqsE or RhlR orthologs were combined at a 1:2 ratio of PqsE to RhlR and 25 μl was saved and combined with 10 μl of lysis buffer and 30 μl of 2x sample buffer for input assessment. Promega MagneHis Ni-particle beads (20 μl per sample) were washed with lysis buffer and re-suspended in lysis buffer at 100 μl per sample, followed by incubation at 4 °C for 1.5 h with inversion. Following incubation, samples were subjected to brief centrifugation at 250 x *g*, placed on a magnetic rack, and the clarified supernatant was aspirated and discarded. Samples were washed 3 times with lysis buffer and 6x-His protein was eluted with two washes of 20 μl elution buffer (20 mM Tris-HCL pH 8.0, 150 mM NaCl, 500 mM imidazole). Eluted protein was mixed 1:1 with 2x sample buffer, boiled at 100°C for 10 min, and 10 μl of each sample was loaded onto SDS-PAGE gels and separated at 35 mA for 35 min. Gels were stained with Coomassie brilliant blue and imaged on a Bio-Rad EZ-doc gel imager.

### Western blot analysis

*P. aeruginosa* cells were cultured from glycerol stocks in 5 mL of LB at 37 °C with shaking at 200 rpm. Overnight cultures were back diluted 1:100 in 20 mL of LB and grown to OD_600_ of ∼2.0 at 37°C with 200 rpm. Cells were pelleted by centrifugation at 10,876 x *g* for 10 min at 4 °C and frozen at −80 °C until lysis. Frozen pellets were resuspended in 1 mL lysis buffer and lysed by sonication (3x 15-s pulses at 25% amplitude). Protein concentration in the whole-cell lysates was quantified using the Pierce BCA protein assay kit (Thermo Fisher Scientific), diluted in 2x sample buffer, and loaded at a 25 mg/mL final protein concentration. Protein was transferred to polyvinylidene difluoride (Bio-Rad) membranes at 80 A for 50 min using a semi-dry transfer cell (Bio-Rad). Following protein transfer, blocking was performed with 1x Tris-buffered saline with Tween 20 (TBST) and 5% milk for 1 h at room temperature with agitation. Primary α-RhlR polyclonal antibody from rabbit (Cambridge Antibodies) was incubated with the membrane at a 1:1,000 dilution for 1 h at room temperature with agitation. Following incubation with the primary antibody, three 5 min washes were performed with TBST before incubation with mouse a-rabbit which was cross adsorbed with secondary antibody conjugated with horseradish peroxidase (Thermo Fisher Scientific) for 1 h at room temperature with agitation at a 1:10,000 dilution. Three 5-minute washes with TBST were performed before the addition of Pierce ECL western blotting substrate (Thermo Fisher Scientific) and imaging on an iBright-1500 (Thermo Fisher Scientific) and quantified using ImageJ software. All antibody solutions were made in 1x TBST and 5% milk.

### Luciferase reporter assays

This luciferase reporter assay was performed as previously described. Briefly, *rhlR* was expressed from the pBAD promoter in an *E. coli* strain containing a p*rhlA*-*luxCDABE* fusion in pCS26. AHL was added to the cultures at the time of induction at a final concentration of 10 µm. The CepR reporter was generated using the same methods, with *cepR* under an arabinose-inducible promoter and a CepR-dependent promoter fused to luciferase (p*cepI-luxCDABE*). For both reporters, assays were also conducted in the presence of *pqsE, pqsE* from *P. fluorescens* NCTC 10783, and *hhqE^Bc^*, or an empty vector control, with expression from the *lac* promoter.

### Catalytic activity and enzyme kinetics

To measure enzyme activity, PqsE or HhqE proteins were purified as described above. Purified protein was diluted to a final concentration of 2 μM in assay buffer (50 mM Tricine, 0.01 % Triton X-100) and added to wells of a 96-well plate in a total volume of 100 μl. MU-butyrate (Sigma) was added to wells at an initial concentration of 8 μm, followed by a series of four 2-fold dilutions. Fluorescence was immediately measured at 30 s intervals for 10 min using a SpectraMax M5 (Molecular Devices) microplate reader, with an excitation wavelength of 360 nm and emission wavelength of 450 nm. Background fluorescence from control wells containing assay buffer only was subtracted at each substrate concentration. Fluorescence intensity at 3 min was used to determine the initial hydrolysis rates to ensure data were on a linear range. Thioesterase activity was assessed in 0.1 M Tris HCl (pH 8.0) buffer with Ellman’s reagent^55^ excess (5,5’-dithiobis(2-nitro-benzoic acid), final concentration 2 mM).The reaction was conducted with acetoacetyl-CoA starting at a maximum concentration of 1 mM followed by four 2-fold serial dilutions. The protein was added to achieve a final concentration of 2 μM in a total assay volume of 100 μl. The reaction progress was monitored by measuring absorbance at 412 nm (ε412 nm = 14,150 M^-1^ cm^-1^) at 30 s intervals for 30 min on a SpectraMax M5 (Molecular Devices) microplate reader. Turnover (*K*_cat_) and the Michaelis-Menten constant (*K*_m_) were determined for PqsE and HhqE proteins using Prism 10.2.3 software.

### Electrophoretic mobility shift assay

HhqE and CepR proteins were diluted in EMSA buffer (200 mM KCl, 50 mM Tris-HCl, 250 mg/mL bovine serum albumin, 50 mM NaCl, 5 nM EDTA, 5 mM MgCl_2_, 5 mM dithiothreitol [pH 8.0]) to a working concentration of 13.9 μM and a series of five 1.5-fold dilutions were made. EMSA reactions comprised 18 μl EMSA buffer, 2 μl protein dilution, and 1 μl of 10 ng/μl *cepI* promoter DNA (300 bp). The reactions were incubated at 30 °C for 15 min. 2 μl of Novex Hi-Density TBE 5 x sample buffer (Thermo Fisher Scientific) was mixed with 8 μl of the EMSA reaction and loaded on an 8.0% agarose gel. Electrophoresis was performed in 0.5 X TB buffer at 100 V for 1 h followed by washing with 0.5 X TB buffer for 15 min. Gels were stained with 50 mL 1 x GelRed in 0.5 X TB buffer for 30 min at RT with shaking, washed three times with 0.5 X TB buffer for 15 min, and visualized on a Bio-Rad EZ-doc gel imager.

### Protein solubility assays

*E. coli* BL21 DE3 (Invitrogen) transformed with pET-DUET containing RhlR or CepR were cultured from glycerol stocks in 3 mL of LB + ampicillin at 37 °C with shaking at 200 rpm. Overnight cultures were back diluted 1:500 in LB + ampicillin (100 μg/mL). Cultures were grown to OD_600_ of 0.5 at 30°C with 220 rpm. For induction, 1 mM IPTG and the desired HSL were added at 150 μM final concentration. Cultures were incubated for 3-4 h at 28 °C with 220 rpm. Cells were harvested at 10,876 x *g*, resuspended in 0.5 mL of lysis buffer (20 mM Tris-HCl pH 7.5, 0.5 mM EDTA, 1 mM DTT), and lysed by sonication (15x 1-s pulses at 50% amplitude). In the case of RhlR, 150 μM was added to the lysis buffer. 50 μl of the whole-cell lysate fraction was collected before isolation of the soluble fraction by centrifugation at 21,300 x *g*. Whole-cell lysate and supernatant fractions were loaded onto an SDS-PAGE gel at a final protein concentration of 50 mg/mL and subjected to electrophoresis at 35 mA for 35 min. Gels were stained with Coomassie Brilliant Blue and imaged on a Bio-Rad EZ-doc gel imager.

### Bacterial two-hybrid assays

Bacterial Adenylate Cyclase Two Hybrid (BACTH) plasmids were constructed using Hifi assembly (NEB) according to the manufacturer’s protocol. Briefly, *abaR* and *blsA* point mutants were constructed using overlap PCR with mutagenic primers and Hifi compatible oligonucleotides (Table S2). Positive clones were sequence verified. pUT18 and pKNT25 plasmids harboring genes of interest were co-transformed into Δ*cyaA E. coli* via electroporation and selected for with ampicillin (100 μg/mL) and kanamycin (50 μg/mL). Transformed Δ*cyaA* strains were grown in selective media overnight and then 3 μl were spotted on dual antibiotic plates with 40 μg/mL X-Gal and top spread with IPTG at a final concentration of 0.5 mM and grown overnight at 30 °C before imaging.

## ACKNOWLEDGMENTS

This work was supported by National Institutes of Health training grant T32GM132066 to C.P.M, NIH grant R01GM14436101, New York Community Trust Foundation grant P19-000454, and Cystic Fibrosis Foundation grant PACZKO21G0 to J.E.P. The authors thank the research laboratories in the Division of Genetics at the Wadsworth Center for helpful discussions on the research and for sharing resources. The authors also thank Dr. Lingyun Li in the Division of Environmental Health Sciences at the Wadsworth Center for mass spectrometry technical expertise, Dr. Izzy Taylor from the College of William and Mary for advice on PqsE kinetic assays, and Dr. Joe Wade in the Division of Genetics at the Wadsworth Center for sharing the Δ*cyaA E. coli* strain. This work was made possible with the help of the dedicated staff scientists at the Advanced Genomics Technologies Center, the Media & Tissue, and the Protein Biochemistry & Immunology Core Facilities at the Wadsworth Center. The authors thank Dr. Karen Chave, Director of the Wadsworth Center Protein Expression and Biochemistry Instrumentation Core Facility, for mass photometry training.

## Code availability

Custom code and all commands issued in this study can be found at https://github.com/calebmallery/PqsE_evolution_manuscript_methods.

## SUPPORTING INFORMATION

**Supplementary FIG 1.**
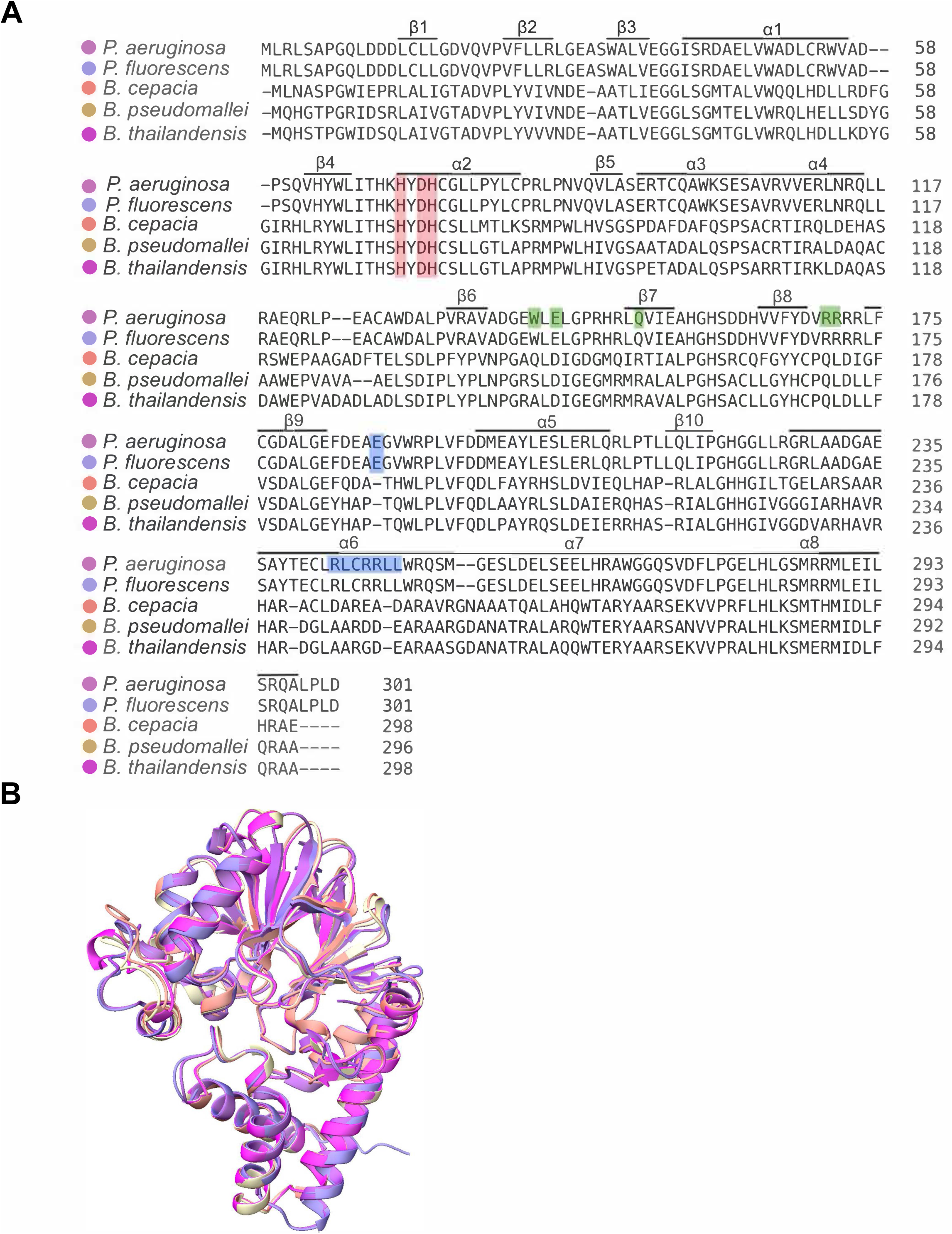
Sequence and structural comparison of PqsE and HhqE orthologs. (a) Multiple-sequence alignment of full-length protein sequences of PqsE from *P. aeruginosa* PA14 and *P. fluorescens* NCTC 10783 or HhqE in *B. cepacia* ATCC 25416, *B. pseudomallei* K96243, and *B. thailandensis* E264. Active site residues H69, H71, D73, and H74 (numbering according to PA14 PqsE) are highlighted in red. Residues involved in PqsE dimerization are highlighted in blue; residues unique to PqsE that are required for the PqsE-RhlR interaction are highlighted in green. Secondary structure assignments are represented above the sequence, determined from the PA14 PqsE experimental structure (PDB: 7KGW^3^). (b) Structural overlay of PA14 PqsE (PDB: 7KGW^3^) with predicted structures of the *Pseudomonas* and *Burkholderia* orthologs. For clarity, structures are depicted as ribbons, with the color scheme corresponding to the alignment in (a). All structure images were generated in ChimeraX^4-6^.

**Supplementary FIG 2.**
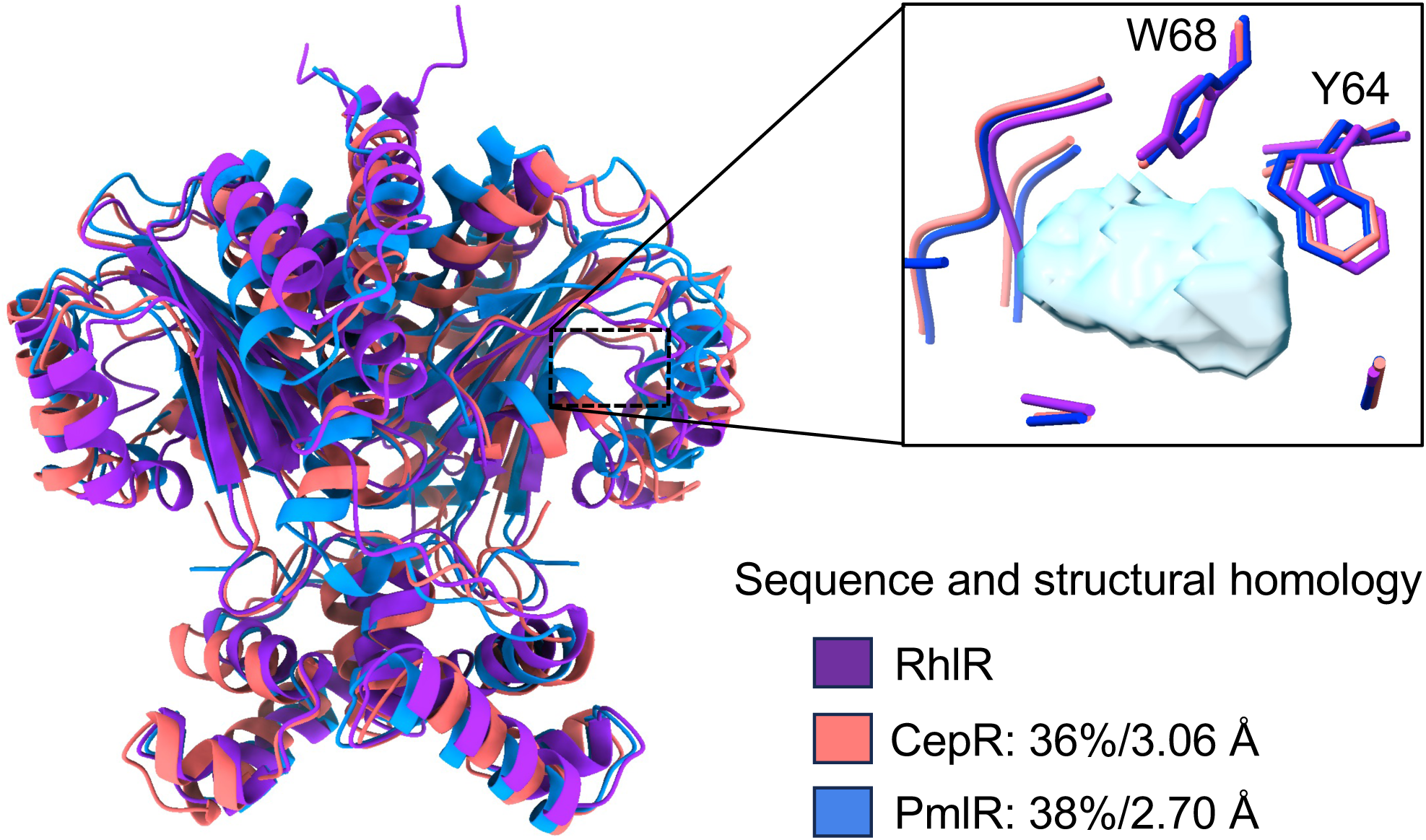
Structural similarities of LuxR-type receptors. Structural overlay of AlphaFold-Multimer predicted structures of CepR from *B. cepacia* and PmlR from *B. pseudomallei* compared to the experimental structure of RhlR (PDB ID: 8DQ0^7^). The percentage amino acid identity and root mean square deviation (rmsd) values of CepR and PmlR relative to PA14 RhlR are indicated. The inset highlights the universally conserved YXXXW motif within the ligand binding pocket (LBP) of LuxR-type receptors (numbering based on PA14 RhlR). The solvent-accessible volume of the RhlR LBP, shown in light blue is included to provide structural context. All structure images were generated in ChimeraX.

**Supplementary FIG 3.**
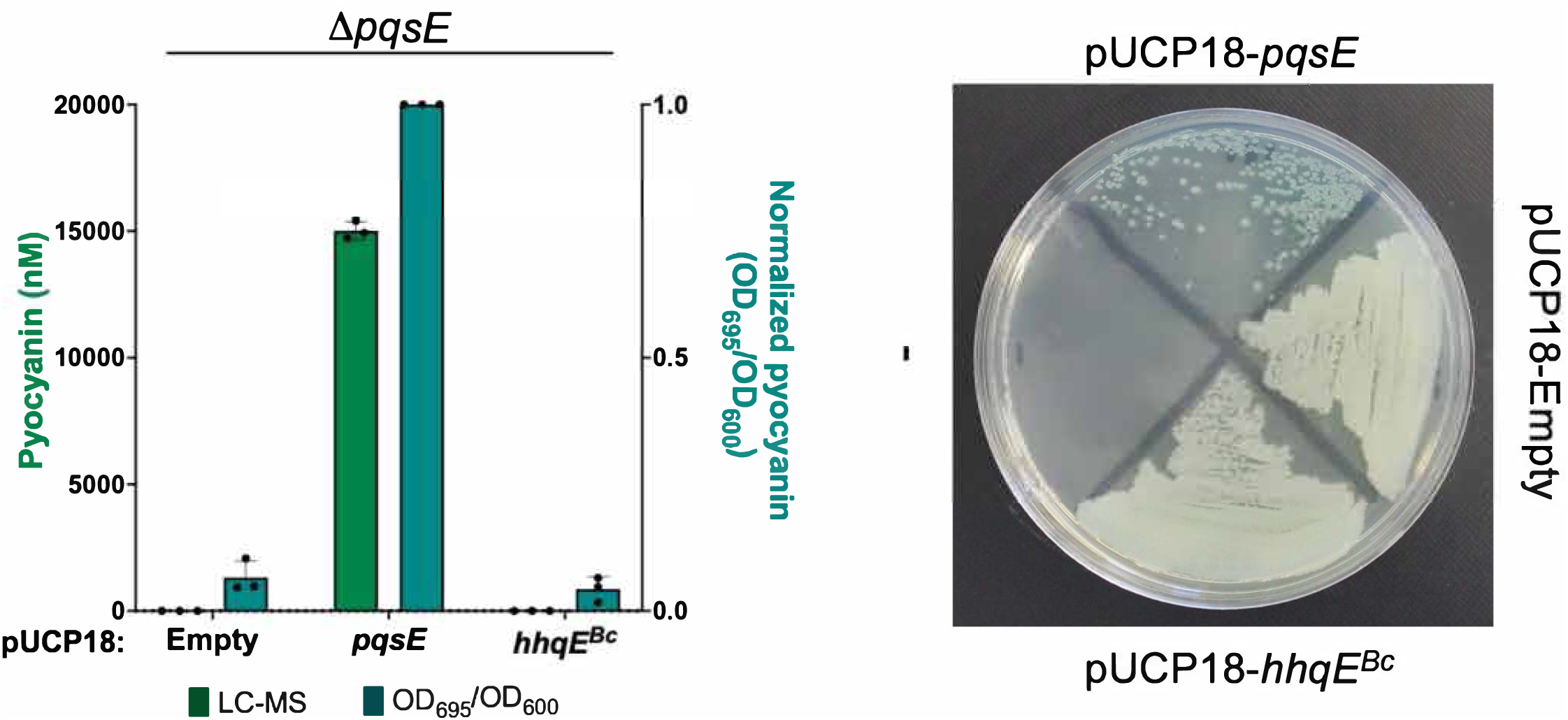
HhqE does not complement pyocyanin production in a Δ*pqsE* strain of *P. aeruginosa*. (a) Pyocyanin production in Δ*pqsE* strains of *P. aeruginosa* expressing plasmid-borne *pqsE* or *hhqE^Bc^* was quantified by OD_695_/OD_600_ and LC-MS. OD_695_/OD_600_ values were normalized to levels observed in the strain expressing WT *pqsE* (center bar). (b) Growth of Δ*pqsE* strains carrying plasmids as in (a) on LB agar. Complementation of pyocyanin production was observed with WT *pqsE* but not with *hhqE* from *B. cepacia*.

**Supplementary FIG 4.**
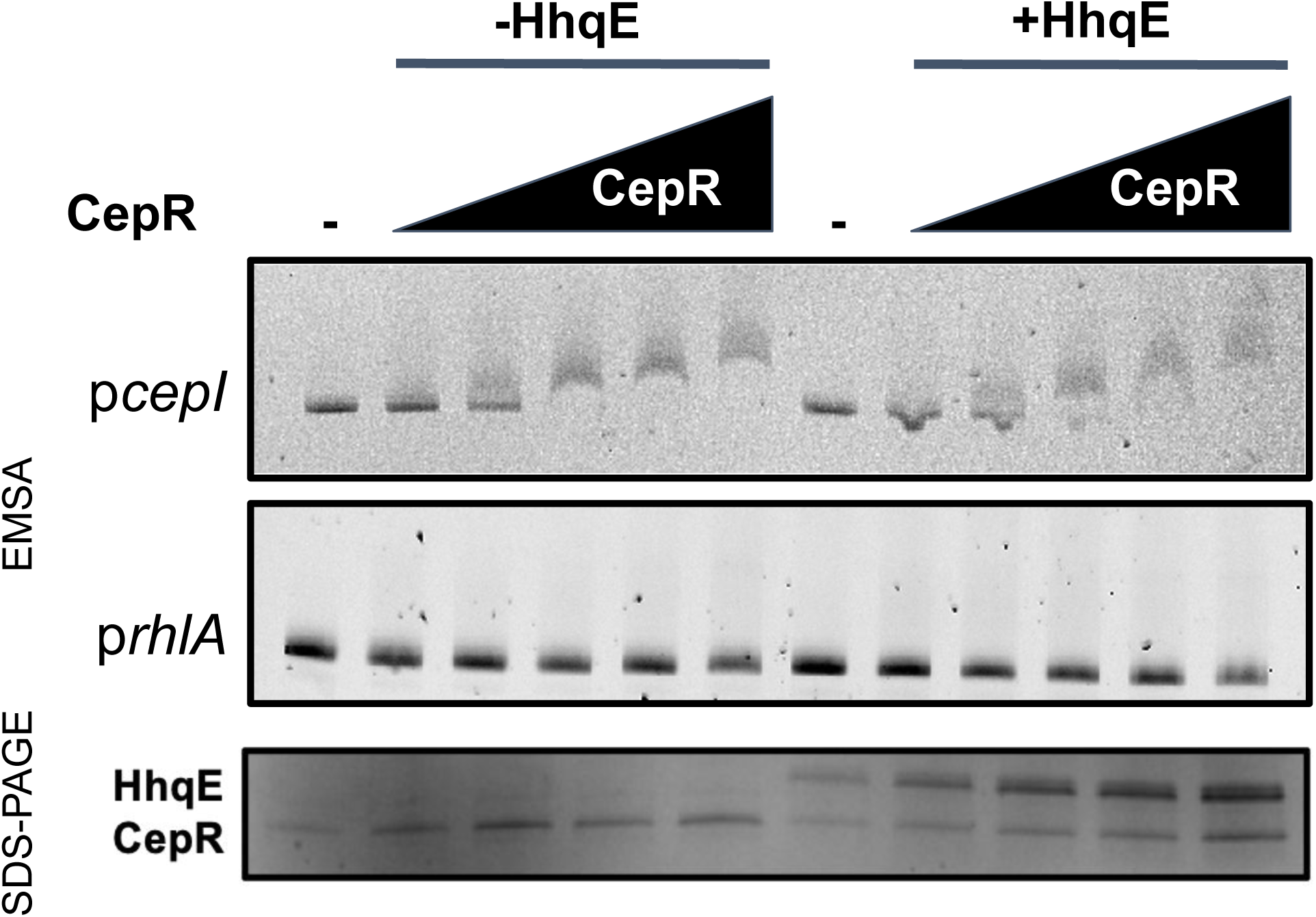
HhqE does not enhance CepR binding to promoter DNA. (a) EMSA analysis of *cepI* promoter DNA alone (minus symbol, left lane), with increasing concentrations of purified CepR:C_8_HSL (left half), and with both CepR and HhqE (right half). An EMSA using *rhlA* promoter DNA was included as a negative control (middle). A representative SDS-PAGE gel shows protein levels for each sample (bottom). Final concentrations of CepR and HhqE were: 0 nm, 280 nM, 420 nM, 630 nM, 950 nM, and 1,350 nM.

**Table S1.**
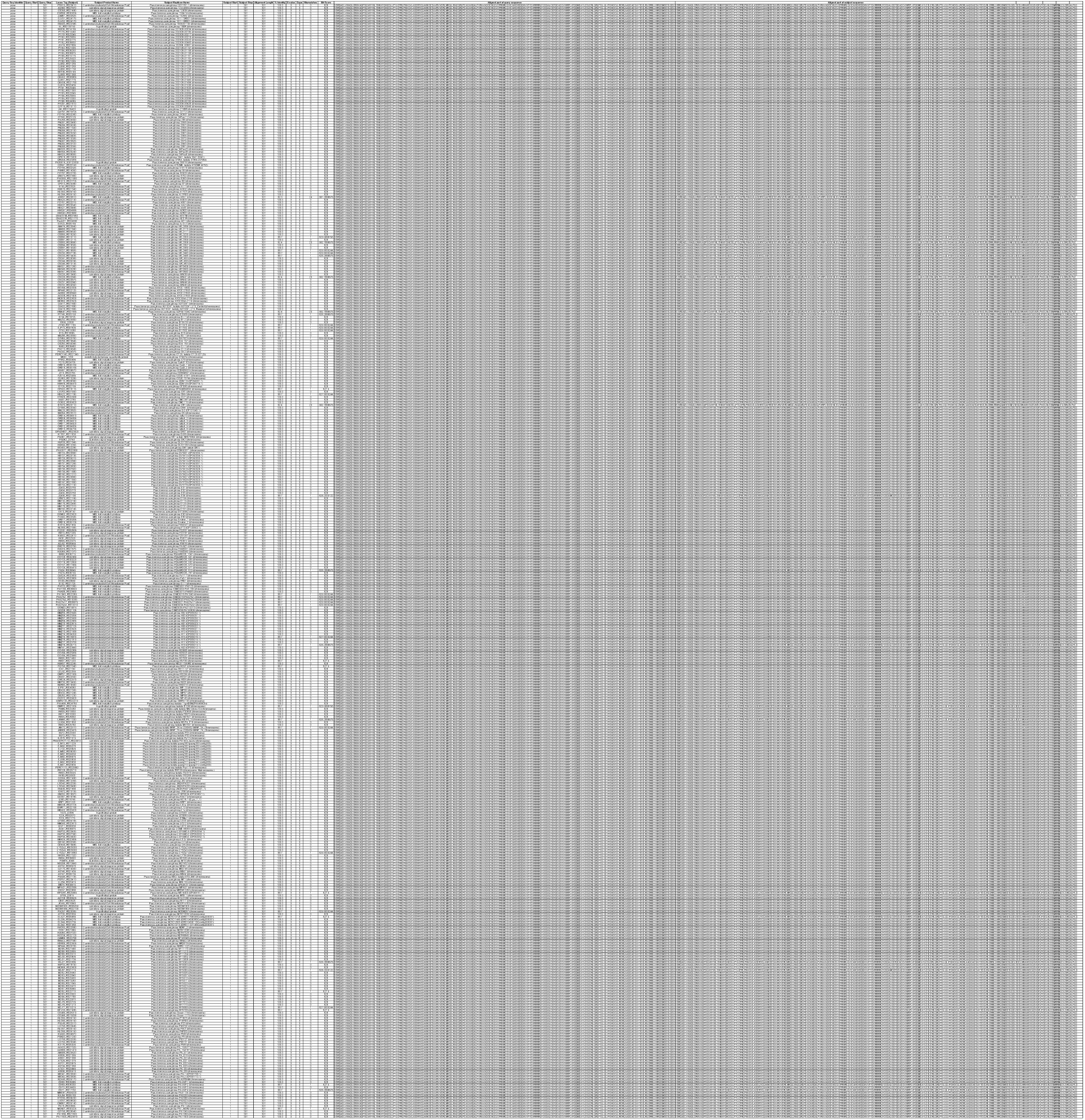
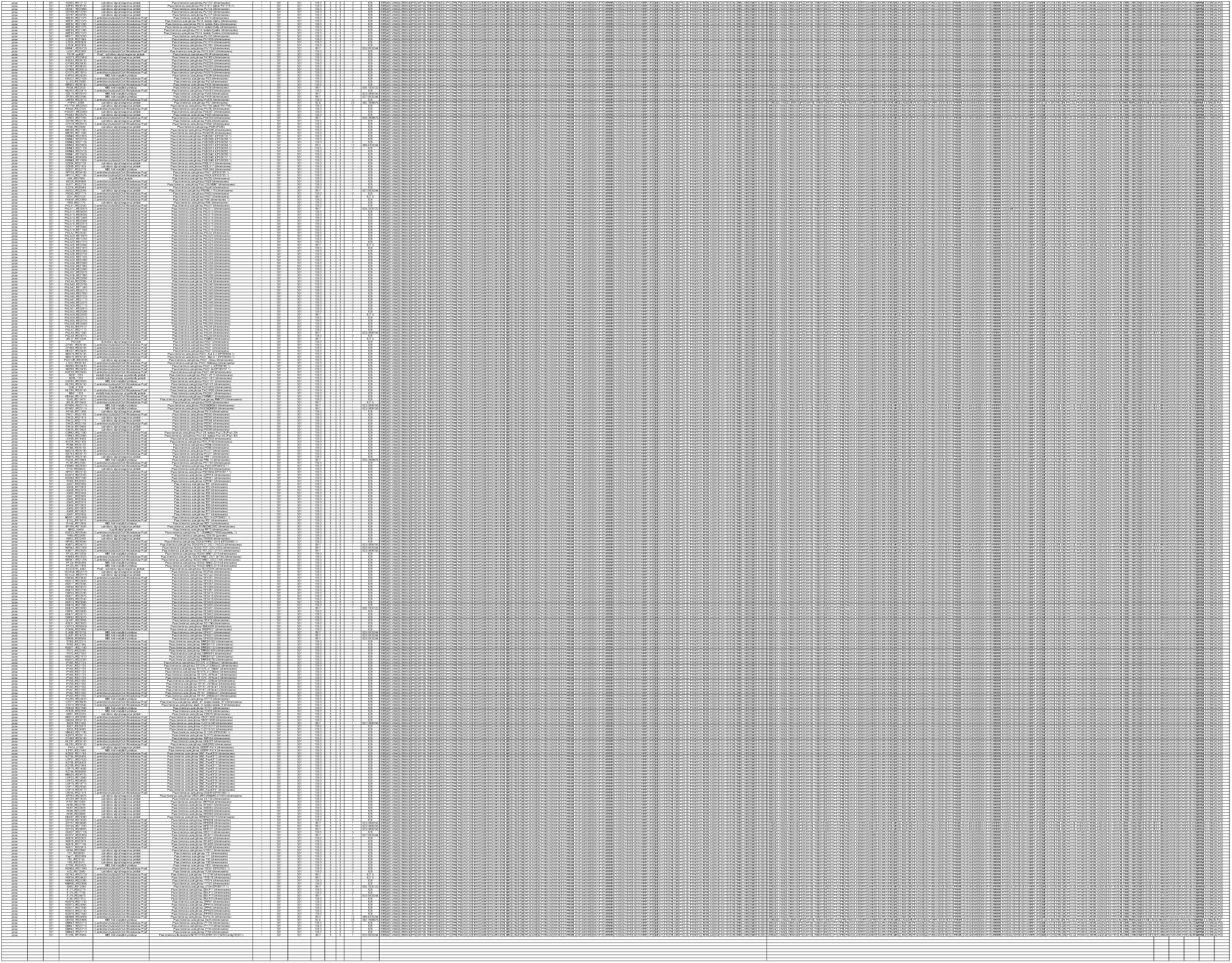
NCBI BlastP results for PqsE homologs obtained from the *Pseudomonas* Genome DB^1^. Included in this table are standard BlastP output with *P. aeruginosa* PqsE used as a reference.

**Table S2.** Structural and sequence alignment table. This table shows the root mean square deviations and TM-score of predicted structures from *P. fluorescens* NCTC 10783, *B. cepacia* ATCC 25416, *B. thailandensis* E264, *B. pseudomallei* K96243 compared to the experimental structure of PqsE from *P. aeruginosa* (PDB ID: 5HIO^2^). This table also includes the percent amino acid identity and protein sequence length of each protein.

| PDB/ID | RMSD | TM-score | Identity | Sequence length |
| --- | --- | --- | --- | --- |
| 5HIO | - | - | - | 301 |
| *P. fluorescens* PqsE | 0.47 | 0.99 | 99% | 301 |
| *B. cepacia* HhqE | 1.80 | 0.92 | 31% | 298 |
| *B. pseudomallei* HhqE | 1.90 | 0.92 | 30% | 298 |
| *B. thailandensis* HhqE | 1.91 | 0.92 | 30% | 298 |

**Table S3.** Strains and plasmids used in this study. This table includes the names, genotypes, antibiotic resistance markers, and sources of all strains used in this study.

| Strain name | Genotype | Plasmid | Antibiotic resistance | Source |
| --- | --- | --- | --- | --- |
| JPS0102 | *E. coli* BL21 | pET28-pqsE | Kan | Mukherjee *et al.* (2018) |
| JPS0103 | *E. coli* BL21 | pET23-rhlR | Amp | McCready *et al.* (2019) |
| JPS0118 | *E. coli* TOP10 | pBAD-A-rhlR pCS26-prhIA-luxCDABE | Amp/Kan | Paczkowski *et al.* (2017) |
| JPS0151 | *P. aeruginosa* ΔrhlR | | | Mukherjee *et al.* (2018) |
| JPS0154 | *P. aeruginosa* ΔrhlI | | | Mukherjee *et al.* (2018) |
| JPS0222 | *P. aeruginosa* (UCBPP-PA14) WT |  |  | Gift from Dr. George O'Toole |
| JPS0225 | *E. coli* DH5a | pET-DUET empty | Amp | Invitrogen |
| JPS0226 | *E. coli* DH5a | pBAD-A empty | Amp | Invitrogen |
| JPS0227 | *E. coli* DH5a | pEXG2 empty | Gent | Gift from Dr. Joseph Mougous |
| JPS0247 | *E. coli* Top 10 | pUCP18 empty | Amp | Mukherjee *et al.* (2018) |
| JPS0268 | *P. aeruginosa* ΔpqsE | pUCP18-empty | Carb | Simanek *et al.* (2022) |
| JPS0278 | *P. aeruginosa* ΔrhlI ΔpqsE | | | Simanek *et al.* (2022) |
| JPS0375 | WT *Burkholderia cepacia* |  |  | ATCC |
| JPS0544 | *P. aeruginosa* PqsE (R243A/R246A/R247A) |  |  | Simanek *et al.* (2022) |
| JPS0561 | *P. aeruginosa* ΔpqsE | | | Simanek *et al.* (2022) |
| JPS0592 | *P. aeruginosa* ΔrhlI PqsE (R243A/R246A/R247A) | | | Simanek *et al.* (2022) |
| JPS0670 | *E. coli* DH5a | pUT18-empty | Amp | This study |
| JPS0671 | *E. coli* DH5a | pUT18-zip | Amp | This study |
| JPS0808 | *E. coli* DH5a | pACYC-empty | Tet | Taylor *et al.* (2021) |
| JPS1067 | *E. coli* DH5a | pET28-hhqE (*B. cepacia*) | Kan | This study |
| JPS1068 | *E. coli* BL21 | pET28-hhqE (*B. cepacia*) | Kan | This study |
| JPS1069 | *E. coli* DH5a | pET-DUET-cepR (*B. cepacia*) | Amp | This study |
| JPS1070 | *E. coli* BL21 | pET-DUET-cepR (*B. cepacia*) | Amp | This study |
| JPS1157 | *E. coli* DH5a | pET28-pqsE chimera (C-term) | Kan | This study |
| JPS1207 | *E. coli* BL21 | pET28-pqsE chimera (C-term) | Kan | This study |
| JPS1223 | *E. coli* DH5a | pET28-pqsE (*P. fluorescens* NCTC 10783) | Kan | This study |
| JPS1224 | *E. coli* DH5a | pET28-hhqE (*B. pseudomallei*) | Kan | This study |
| JPS1225 | *E. coli* DH5a | pET28-pqsE (*P. fluorescens* NCTC 10783) | Kan | This study |
| JPS1228 | *E. coli* DH5a | pET-DUET-pmlR (*B. pseudomallei*) | Amp | This study |
| JPS1233 | *E. coli* BL21 | pET28-pqsE (*P. fluorescens* NCTC 10783) | Kan | This study |
| JPS1234 | *E. coli* BL21 | pET-DUET-pmlR (*B. pseudomallei*) | Amp | This study |
| JPS1305 | *E. coli* DH5a | pUCP18-hhqE (*B. cepacia*) | Amp | This study |
| JPS1308 | *P. aeruginosa* ΔpqsE | pUCP18-hhqE (*B. cepacia*) | Carb | This study |
| JPS1470 | *E. coli* DH5a | pKNT25 | Kan | Gift from Dr. George O'Toole |
| JPS1471 | *E. coli* DH5a | pKNT25-zip | Kan | Gift from Dr. George O'Toole |
| JPS1475 | *E. coli* Top 10 | pKNT25-abaR | Kan | This study |
| JPS1511 | *E. coli* DH5a | pBAD-A-cepR (*B. cepacia*) | Amp | This study |
| JPS1547 | *E. coli* Top 10 | pUT18-blsA | Amp | This study |
| JPS1555 | *E. coli* MG1655 ΔcyaA | pKNT25-abaR/pUT18-blsA | Amp/Kan | This study |
| JPS1556 | *E. coli* MG1655 ΔcyaA | pKNT25-abaR/pUT18 | Amp/Kan | This study |
| JPS1557 | *E. coli* MG1655 ΔcyaA | pKNT25/pUT18-blsA | Amp/Kan | This study |
| JPS1558 | *E. coli* MG1655 ΔcyaA | pKNT25/pUT18 zip | Amp/Kan | This study |
| JPS1570 | *E. coli* Top 10 | pUT18-blsA R99A | Amp | This study |
| JPS1584 | *E. coli* MG1655 ΔcyaA | pUT18-blsA R99A/pKNT25-abaR | Amp/Kan | This study |
| JPS1586 | *E. coli* MG1655 ΔcyaA | pUT18-blsA R99A/pKNT25 | Amp/Kan | This study |
| JPS1593 | *E. coli* Top 10 | pACYC-hhqE (*B. cepacia*) | Tet | This study |
| JPS1646 | *E. coli* Top 10 | pBAD-A-cepR pCS26-pcepI-luxCDABE | Amp/Kan | This study |
| JPS1647 | *E. coli* Top 10 | pBAD-A-rhlR pCS26-prhIA-luxCDABE pACYC-hhqE (*B. cepacia*) | Amp/Kan/Tet | This study |
| JPS1661 | *E. coli* DH5a | pKNT25-abaR E6A/D7A | Kan | This study |
| JPS1663 | *E. coli* Top 10 | pBAD-A-empty pCS26-pcepI-luxCDABE | Amp/Kan | This study |
| JPS1669 | *E. coli* MG1655 ΔcyaA | pUT18-blsA/pKNT25-abaR E6A/D7A | Amp/Kan | This study |
| JPS1670 | *E. coli* MG1655 ΔcyaA | pUT18/pKNT25-abaR E6A/D7A | Amp/Kan | This study |
| JPS1715 | *E. coli* Top 10 | pACYC-pqsE (*P. fluorescens* NCTC 10783) | Tet | This study |
| JPS1732 | *E. coli* Top 10 | pBAD-A-rhlR pCS26-prhIA-luxCDABE pACYC-pqsE (*P. fluorescens* NCTC 10783) | Amp/Kan/Tet | This study |
| JPS1737 | *E. coli* DH5a | pUCP18-pqsE (*P. fluorescens* NCTC 10783) | Amp | This study |
| JPS1918 | *E. coli* DH5a | pEXG2-lon | Gent | This study |
| JPS1919 | *E. coli* SM10-λpir | pEXG2-lon | Gent | This study |
| JPS1920 | *P. aeruginosa* Δlon | | | This study |
| JPS1921 | *P. aeruginosa* ΔpqsE/Δlon | | | This study |

**Table S4.**
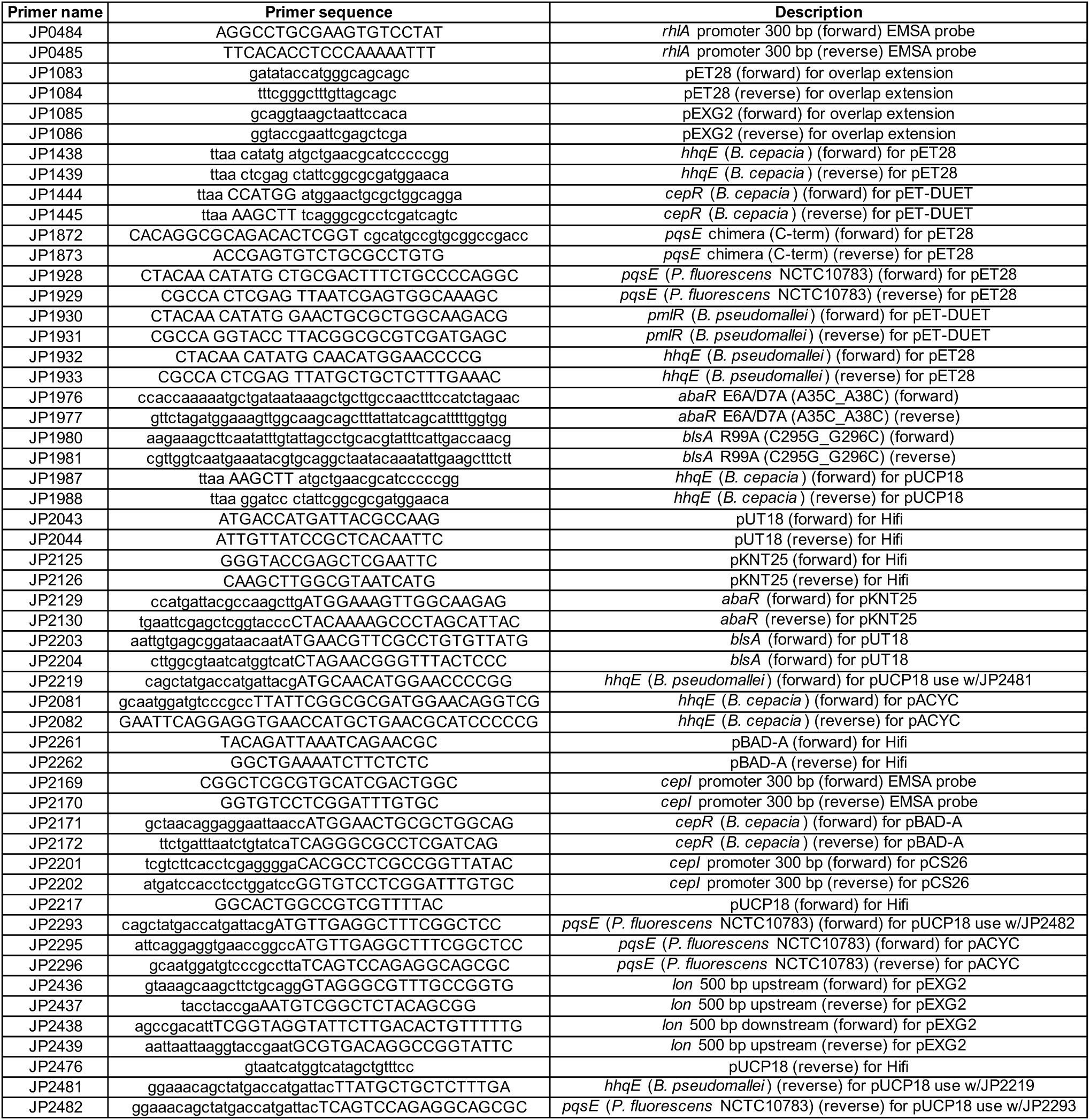
Oligonucleotides used in this study. This table includes the names, sequences (5’ to 3’), and description of each oligonucleotide used in this study.

